# Evolution of enhanced innate immune suppression by SARS-CoV-2 Omicron subvariants

**DOI:** 10.1101/2022.07.12.499603

**Authors:** Ann-Kathrin Reuschl, Lucy G. Thorne, Matthew V.X. Whelan, Roberta Ragazzini, Wilhelm Furnon, Vanessa M. Cowton, Giuditta de Lorenzo, Dejan Mesner, Jane L. E. Turner, Giulia Dowgier, Nathasha Bogoda, Paola Bonfanti, Massimo Palmarini, Arvind H. Patel, Clare Jolly, Greg. J. Towers

## Abstract

SARS-CoV-2 adaptation to humans is evidenced by the emergence of variants of concern (VOCs) with distinct genotypes and phenotypes that facilitate immune escape and enhance transmission frequency. Most recently Omicron subvariants have emerged with heavily mutated spike proteins which facilitate re-infection of immune populations through extensive antibody escape driving replacement of previously-dominant VOCs Alpha and Delta. Interestingly, Omicron is the first VOC to produce distinct subvariants. Here, we demonstrate that later Omicron subvariants, particularly BA.4 and BA.5, have evolved an enhanced capacity to suppress human innate immunity when compared to earliest subvariants BA.1 and BA.2. We find that, like previously dominant VOCs, later Omicron subvariants tend to increase expression of viral innate immune antagonists Orf6 and nucleocapsid. We show Orf6 to be a key contributor to enhanced innate immune suppression during epithelial replication by BA.5 and Alpha, reducing innate immune signaling through IRF3 and STAT1. Convergent VOC evolution of enhanced innate immune antagonist expression suggests common pathways of adaptation to humans and links VOC, and in particular Omicron subvariant, dominance to improved innate immune evasion.

---

Alpha, Delta and Omicron have been sequentially globally dominant, with each VOC evolving independently from early lineage SARS-CoV-2 virus. Since the appearance of the Omicron lineage, Omicron subvariants have evolved, co-circulated, recombined and replaced each other locally or globally. Following the first dominant Omicron subvariants BA.1 and BA.2, BA.4 and BA.5 emerged with each displaying increasing levels of antibody escape, through mutation of spike, threatening vaccine efficacy and increasing hospitalisations ^1,2, 3–14^. In addition, Omicron subvariants are accumulating mutations across several of the genome, consistent with ongoing adaptation to host.

To understand phenotypic differences between Omicron subvariants, and the selective forces driving their evolution, we compared replication of, and host responses to, BA.1-BA.5 with Delta, the previously dominant VOC, in Calu-3 human airway epithelial cells (Fig. 1). We equalized input dose of each variant by viral envelope (E) gene copies (RT-qPCR) as this ensures cells are exposed to equal starting amounts of viral RNA, which is the major viral PAMP activating defensive host innate immune responses^15,16^. Most importantly, this approach normalizes dose independently of variant-specific differences in cell tropism or entry routes (Fig. 1a, Extended Data Fig. 1a,b)^17–19^, which we and others have shown impact both titer determination and input equalization by cell-line infectivity measurements such as TCID50 or plaque assay (Extended Data Fig. 1d-f). Our approach is particularly relevant for comparing Omicron subvariants because spike mutations have been shown to alter tropism, increasing cathepsin-dependent endosomal entry and reducing dependence on cell surface TMPRSS2^17,18,20,21^, irrespective of virion spike cleavage efficiency (Extended Data Fig. 1c). Endosomal cathepsins or cell surface TMPRSS2 are required to cleave spike prior to ACE2 mediated entry^22,23^. Indeed, in line with previously published data^17,19^ we have found that Omicron, particularly BA.5, has enhanced entry (cathepsin dependent and E64d-sensitive) in TMPRSS2-negative cells such as Hela-ACE2 compared to previous VOCs such as Delta, whereas entry into Calu-3 cells is largely TMPRSS2-dependent (camostat-sensitive) (Extended Data Fig. 1a.b) resulting in striking cell-type specific differences between variant titers by TCID50 (Extended Data Fig. 1f).

**Fig. 1.**
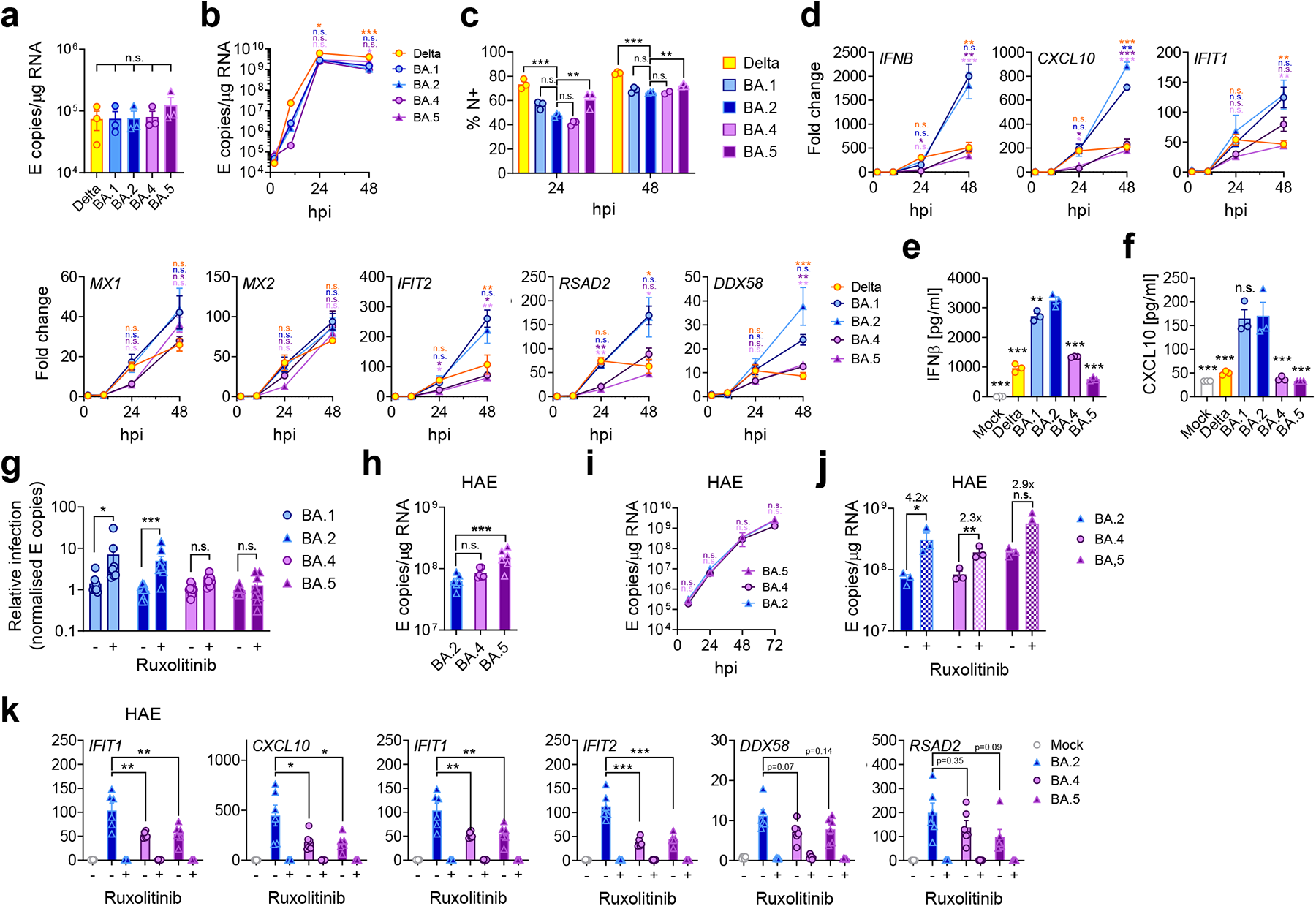
BA.5 displays enhanced innate immune antagonism during infection of airway epithelial cells. **(a-f)** Calu-3 infection with 2000 E copies/cell of Delta (yellow; Ο), BA.1 (blue; Ο), BA.2 (blue; Δ), BA.4 (purple; Ο) and BA.5 (purple; Δ), n=3. (**a**) Mean viral E copies at 2hpi across 3 independent experiments at 2000 E copies/cell. (**b**) Viral replication over time measured by RT-qPCR for intracellular E copies. (**c**) Infection levels measured by nucleocapsid expression (% N+ by flow cytometry). (**d**) Expression of *IFNB*, *CXCL10*, *IFIT1, IFIT2, RSAD2, MX1, MX2* and *DDX58* in infected cells over time. (**e**) IFNβ and (**f**) CXCL10 secretion from infected Calu-3 cells measured by ELISA at 48hpi. (**g**) Rescue of viral replication by JAK1-inhibitor ruxolitinib in Calu-3 cells at 48hpi at 2000 E copies/cell. Shown are the relative infection levels across three independent experiments determined by E copies/μg RNA normalized to the median infection level of the untreated control. (h-k) Primary bronchial human airway epithelial cells (HAEs) were infected with 1500 E copies/cell of the indicated variants, n=3. Viral replication was measured by (**h**) intracellular E copies at 72hpi and (**i**) viral release into apical washes over time. (**j**) Intracellular viral E copies in HAEs in the presence or absence of ruxolitinib at 72hpi. (**k**) Expression of *IFNB*, *CXCL10, IFIT1*, *IFIT2*, *DDX58* and *RSAD2* in cells from (j). Fold changes are normalized to mock at (d) 2hpi or (k) 72hpi. For statistical comparisons one-way ANOVA with Dunnett’s post-test was used to compare all variants at 2hpi in (a) or to compare BA.2 with other variants at 24 and 48hpi respectively (b-f). Colors indicate comparator (Delta, yellow; BA.1, blue; BA.4, purple; BA.5, pink). For g and j, indicated comparisons were performed using an unpaired one-tailed Student’s t-Test. For h and k, indicated comparisons were performed using one-way ANOVA with Dunnett’s post-test. For i, Two-way ANOVA with Bonferroni post-test were used to compare variants against BA.2. Mean+/-SEM or individual datapoints are shown. hpi, hours post infection. *, p<0.05; **, p<0.01; ***, p<0.001; n.s., not significant or exact p-value given (k).

Infection of Calu-3 cells with 2000 E gene copies/cell (Fig. 1) or 200 E copies/cell (Extended Data Fig. 2) gave comparable E RNA (RT-qPCR) at 2 hours post infection (hpi), consistent with equal input doses (Fig. 1a and Extended Data Fig. 2i). E gene measurements during infection revealed that Omicron isolates BA.1, BA.2, BA.4 and BA.5 replicated similarly, lagging behind Delta in Calu-3 cells (Fig. 1b, Extended Data Fig. 1k-n and Extended Data Fig. 2h). BA.4 replicated most slowly initially but caught up with BA.1, BA.2 and BA.5 by 24 hpi (Fig. 1b, Extended Data Fig. 1 and Extended Data Fig. 2h). Importantly, these replication differences were observed consistently across several experiments (Fig. 1, Extended Data Fig. 1 and Extended Data Fig. 2). As E gene measurement during infection captures gRNA as well as E, S and Orf3 sgRNAs, we compared the levels of intracellular E RNA with those of Nsp12 and Orf1a (Compare Extended Data Fig. 1k-l to Fig. 1b and Extended Data Fig. 1m-n to Extended Data Fig. 2h), which are uniquely encoded within gRNA. Importantly, the ratio of E to Nsp12 was similar until 24hpi reflecting equivalent levels of E sgRNA synthesis between variants (Extended Data Fig. 1o). Quantification of released virions by measuring E and Nsp12 RNA copies in the supernatant mirrored viral replication (Extended Data Fig 1g-i). Similar patterns of infection were also seen when quantified by intracellular Nucleocapsid (N) staining (Fig. 1c and Extended Data Fig. 2j).

We next compared the host innate immune response to Omicron subvariants during infection of Calu-3 cells. All viral stocks were prepared in human gastrointestinal Caco-2 cells as they are naturally permissive to SARS-CoV-2 replication but do not mount a strong innate response to this infection^15,24^. We confirmed that viral stocks prepared in Caco-2 cells (the highest viral inoculum for each variant was 2000 E copies/cell) did not contain measurable IFNβ and negligible IFNλ1/3 (ELISA) (Extended Data Fig 1p,q), ensuring differences in innate immune activation in Calu-3 infections were not a result of IFN carryover in the viral stocks.

Strikingly, we found that infection of Calu-3 cells with BA.4 and BA.5 resulted in significantly less innate immune activation compared to BA.1/BA.2, evidenced by lower induction of Interferon-β (*IFNB*) and interferon stimulated genes (ISGs) including inflammatory chemokine *CXCL10* and *RSAD2*, *DDX58*, *IFIT1* and *IFIT2* (Fig. 1d and Extended Data Fig 2c,d,l,m) and a trend towards reduced *MX1* and *MX2* expression (Fig. 1d). Reduced host responses to BA.4 and BA.5 infection were also evident at the level of IFNβ and CXCL10 secretion (Fig.1e,f). Slower replication of BA.4 likely contributes in part to the lesser innate immune activation during Calu-3 infection, but BA.5 replication was similar to BA.1 and BA.2 and nonetheless induced significantly less innate immune responses. Inhibition of IFN-mediated JAK/STAT signaling with ruxolitinib, evidenced by the absence of ISG induction (Extended Data Fig. 2d,l), rescued BA.1 and BA.2 infection in Calu-3 cells to a greater degree than BA.4 or BA.5 (Fig. 1g and Extended Data Fig. 2b,n), suggesting that the greater induction of IFNβ by BA.1 and BA.2 reduced their infectivity. BA.1-5 showed similar sensitivities to a range of IFN doses used to pre-treat Calu-3 cells (Extended Data Fig. 2e-g). We therefore conclude that the differences in ruxolitinib sensitivity reflect differences in IFN induction after Calu-3 infection and not differences in IFN sensitivity. Infecting Calu-3 cells with lower virus input doses (200 E copies/cell) recapitulated our observation that Delta replicated better than Omicron BA.1-BA.5 (Extended Data Fig. 2h-j), and we again saw reduced innate immune activation by BA.4 and BA.5 compared to BA.1 and BA.2 (Extended Data Fig. 2l,m). At this lower inoculum, BA.4 infectivity was also strongly rescued by ruxolitinib treatment consistent with its slower replication being due to IFN induction (Extended Data Fig. 2k).

We next compared Omicron subvariant replication and host responses in primary human airway epithelial (HAE) cultures, which better recapitulate the heterogenous polarized epithelial layer of the respiratory tract. We have previously reported that HAEs reveal differences in VOC replication that likely reflect host adaptation, which are not always apparent in highly-permissive cell lines, such as Calu-3^16,25^. Concordantly, BA.5 viral replication was higher than BA.2 and BA.4 in differentiated primary bronchial HAEs at 72hpi, while apical viral release over time was comparable (Fig. 1h,i). Despite BA.4 and BA.5 replicating similarly to parental BA.2 in HAEs, we consistently observed reduced innate activation, measured by ISG induction, after BA.4 and BA.5 infection (*IFNB, CXCL10, IFIT1, IFIT2, DDX58* and *RSAD2*) (Fig. 1k). Inhibiting IFN signaling with JAK-inhibitor ruxolitinib suppressed ISG induction (Fig. 1k) and rescued replication of BA.2 to a greater degree than BA.4 and BA.5 (Fig. 1j). Altogether, data in Figure 1 suggest adaptation to reduce innate immune activation between the earliest (BA.1, BA.2) and subsequent (BA.4, BA.5) Omicron subvariants.

SARS-CoV-2, and other respiratory viruses, reportedly replicate more efficiently in nasal and tracheal epithelial cells^26^, in part due to reduced innate activation and interferon-responsiveness at the lower temperatures of the upper airway^27–29^. To investigate whether lower temperatures reveal further Omicron subvariant adaptation, we compared replication at 32°C in Calu-3 cells. We found BA1-5 all replicated less well than at 37°C (Extended Data Fig. 3a,b) whereas Delta replication was not as temperature-sensitive. As expected^28^, innate immune activation in response to infection, or to RNA sensing agonist poly(I:C), was largely abolished at 32°C (measured by *IFNB* and *CXCL10* mRNA induction) (Extended Data Fig. 3c-e). At 37°C, we again observed lower innate activation for BA.4 and BA.5 compared to BA.1/BA.2. In HAE, lowering the temperature to 32°C did not impact viral replication to the same extent as in Calu-3 cells (Extended Data Fig. 3f). However, we observed reduced virus output in apical washes from infected HAE cultures for all Omicron isolates (Extended Data Fig. 3g-i). Infected HAEs at 32°C also expressed significantly less *IFNB* and *CXCL10* (Extended Data Fig. 3j,k). Overall, our data suggest that Omicron does not replicate better at 32°C in lung epithelial cells even in the absence of an innate immune response. However, it is possible that the intra-tissue temperature throughout the airways remains closer to 37°C than the exhaled breath temperature of 32°C suggests^30^.

We next investigated the mechanism underlying differential innate immune activation by Omicron subvariants. IRF3 and STAT1 are key transcription factors responding to intracellular RNA sensing, exemplified here by poly(I:C) treatment (Extended Data Fig. 4a and Fig. 3w,x). We and others have shown SARS-CoV-2 activates transcription factors IRF3 and STAT1 downstream of RNA sensing ^15,31^. Consistent with their reduced innate immune triggering, we found Omicron BA.4 and BA.5 infection activated significantly less IRF3 phosphorylation than BA.2 infection (Fig. 2a-c). A similar trend was observed for STAT1 serine 727 phosphorylation which is essential for full STAT1 transcriptional activity^32^, but not upstream JAK1-dependent tyrosine 701-phosphorylation (Fig. 2a,d-f). Reduction of STAT1-phosphorylation correlated with reduced STAT1 nuclear translocation during BA.4 and BA.5 infection compared to BA.2, measured by high-content single-cell immunofluorescence imaging of infected nucleocapsid-positive Calu-3 cells (Fig. 2g). These data suggest BA.4 and BA.5 more effectively prevent intracellular activation of innate sensing pathways. We previously reported that SARS-CoV-2 VOC Alpha evolved enhanced innate immune evasion by increasing expression of key innate antagonists Orf6, Orf9b and N (Extended Data Fig. 4m), which manipulate host cell innate immune pathways^16^. To investigate whether Omicron subvariants have also independently evolved enhanced innate immune suppression through similar mechanisms during human adaptation, we measured viral innate antagonist protein expression during infection. Strikingly, we found that BA.4, and particularly BA.5, expressed higher levels of Orf6 and N compared to BA.1 and BA.2 (Fig. 2i-m and Extended Data Fig. 4b-h,j), measured at 48hpi in Calu-3 cells when E RNA levels were equivalent (Fig. 2h). Unlike previous VOCs^16,25^, expression of innate immune antagonist Orf9b was not detected for any Omicron isolate, possibly due to Omicron subvariants encoding lineage-specific Orf9b mutations (P10S and ΔENA at positions 27-29) altering antibody binding and precluding detection by immunoblot (Fig. 2r and Extended Data Fig. 4m). Importantly, Orf9b remained readily detectable in Delta-infected cells (Fig. 2r). Upregulation of Orf6 and N expression by BA.5 was validated using a second independent isolate (Extended Data Fig. 4j-l), and was also evident in lysates from infected HAEs (Extended Data Fig. 4i). Blocking IFN signaling with ruxolitinib rescued replication of all Omicron isolates as before (Fig. 1 and Extended Data Fig. 2) and enhanced viral protein detection by immunoblot (Fig. 2i,r and Extended Data Fig. 4b). Importantly, higher levels of BA.4 and BA.5 Orf6 and N remained apparent after ruxolitinib treatment (Fig. 2i,l,m). We previously showed enhanced levels of Orf6, N and Orf9b protein by Alpha were associated with increased levels of the corresponding sgRNAs^16^. By contrast, BA.5 Orf6 and N sgRNA levels (normalized to genomic Orf1a) were not enhanced, and were only slightly upregulated during BA.4 infection (Fig. 2n,o), particularly in comparison to Alpha (Extended Data Fig. 4n-p). No differences were observed in S and Orf3a sgRNAs which served as controls to rule out a general enhancement of sgRNA synthesis (Fig. 2p,q). Although Omicron subvariants have synonymous and non-synonymous mutations in Orf6 and N, there are no mutations that distinguish BA.4 and BA.5 from BA.1 and BA.2 that provide a simple explanation for increased Orf6 or N protein levels, including in their transcriptional regulatory sequences (TRS) (Fig. 1 and 2, Extended Data Table 1 and 2). Thus, we hypothesize that BA.4 and BA.5 have either evolved independent, novel mechanisms to increase Orf6 and N protein levels, or that the increase is mediated by changes elsewhere in the genome, which may impact viral translation or protein stability. Further studies are required to pinpoint the adaptations regulating Orf6 and N expression levels.

**Fig. 2.**
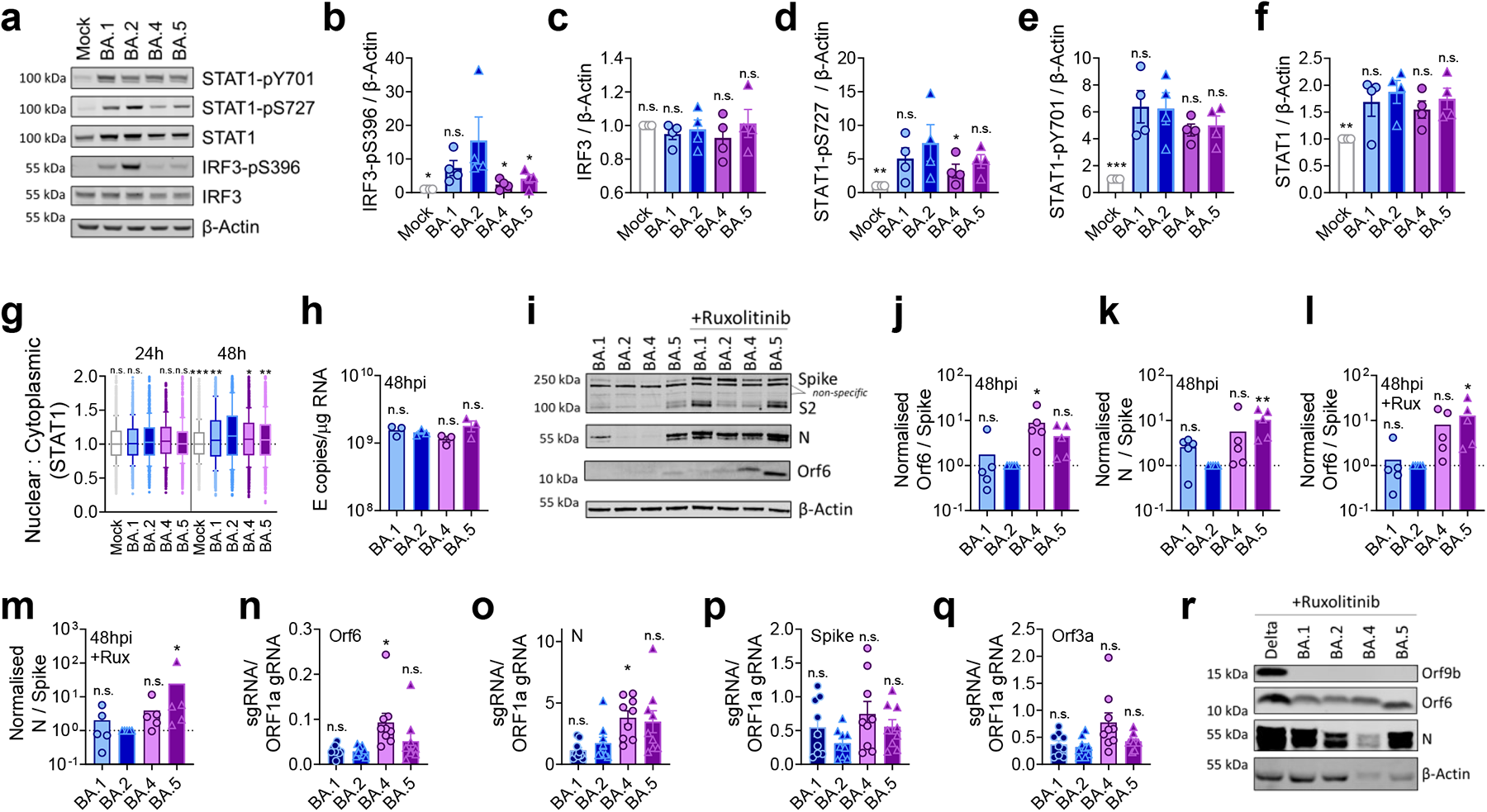
BA.5 efficiently expresses SARS-CoV-2 innate antagonists during airway epithelial cell infection. Calu-3 cells were infected with 2000 E copies/cell of the indicated variants. (**a**) Western blot of STAT1-pY701, STAT1-pS727, total STAT1, IRF3-pS396, total IRF3, and β-Actin at 24hpi. (**b-f**) Quantification of four independent western blots showing (**b**) P-IRF3, (**c**) IRF3, (**d**) STAT1-pS727 (**e**) STAT1-pY701 and (**f**) STAT1 over β-Actin at 24hpi. (**g**) Quantification of STAT1 translocation detected by single-cell fluorescence microscopy over time in Calu-3 cells infected with the indicated variants at 2000 E copies/cell. In infected cultures, translocation was determined in N+ cells. Data from 1500 cells/condition are shown. (**h**) Viral replication at 48hpi. (**i**) Representative western blot of Orf6, N, spike/S2 and β-Actin at 48hpi in infected cells +/-5μM ruxolitinib. Non-specific bands detected by polyclonal anti-spike primary antibody are indicated (see Extended Data Fig. 2a for Mock). (**j-m**) Quantification of Orf6 and N expression from five independent western blots of Calu-3 cells in the absence (**j**, Orf6; **k**, N) or presence of 5μM ruxolitinib (**l**, Orf6; **m**, N), normalized to spike over BA.2. (**n-q**) sgRNA expression of (**n**) Orf6, (**o**) N, (**p**) spike and (**q**) Orf3a normalized to Orf1a genomic RNA in Calu-3 cells at 48hpi, n=9. (**r**) Western blot of Calu-3 cells infected with Delta, BA.1, BA.2, BA.4 and BA.5 at 2000 E copies/cell showing Orf9b, Orf6, N and β-Actin expression at 48hpi+5μM ruxolitinib. For b-f, h, j-q, one-way ANOVA with Dunnett’s post-test was used to compare BA.2 with other variants. For g Kruskal-Wallis test was used to compare groups at 24hpi and 48hpi. Mean+/-SEM or individual datapoints are shown. hpi, hours post infection. sgRNA, subgenomic RNA. *, p<0.05; ***, p<0.001; n.s., not significant.

**Fig. 3.**
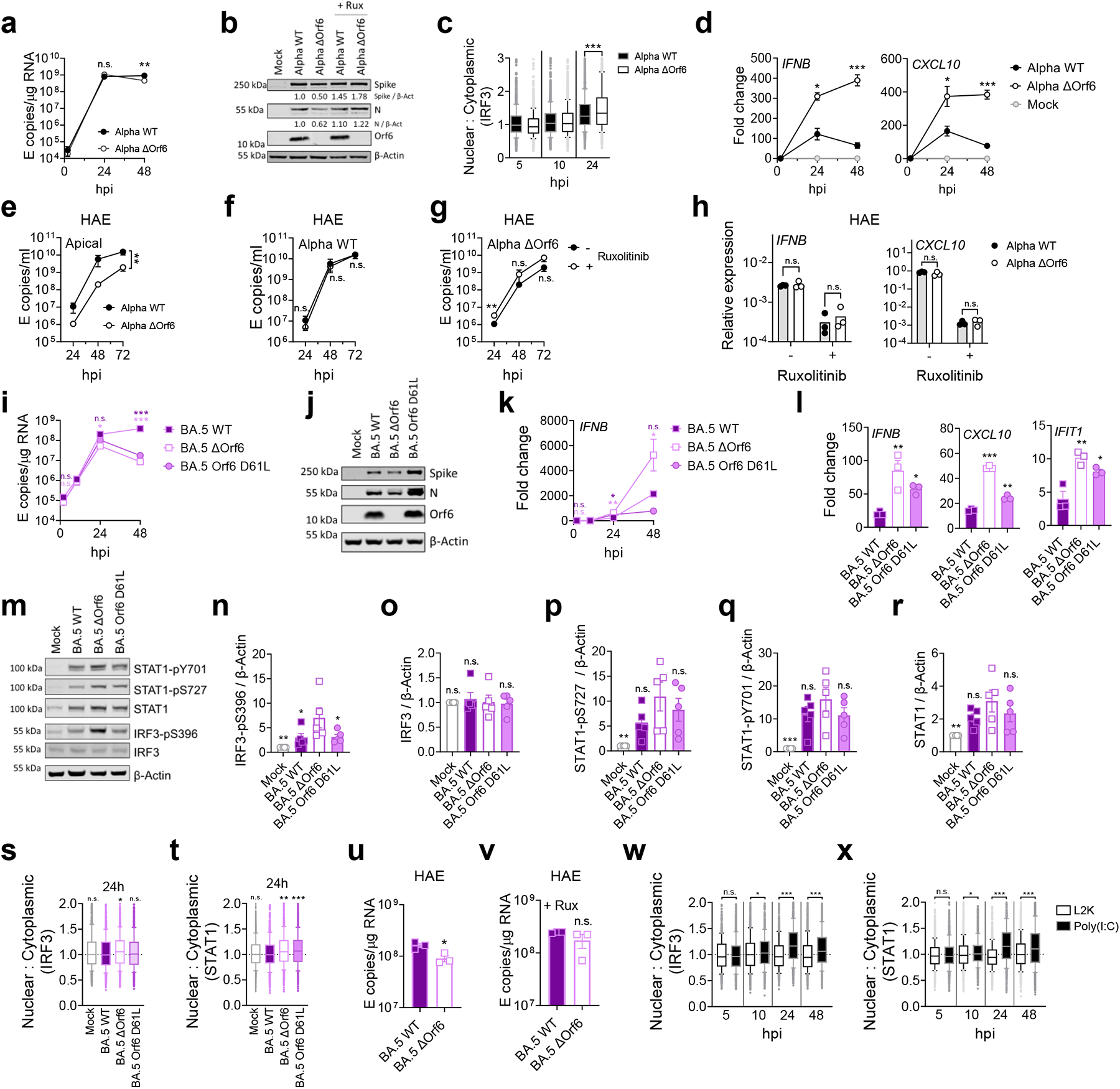
Orf6 expression is a major determinant of enhanced innate immune antagonism by emerging VOCs. (**a**) Replication of reverse genetic (RG) viruses parental Alpha WT and ΔOrf6 in Calu-3 cells infected with 2000 E copies/cell over time, n=3. (**b**) Western blot of RG virus infections in Calu-3 cells at 24hpi for spike, N, Orf6 and β-Actin +/-5μM ruxolitinib. (**c**) Quantification of IRF3 translocation detected by single-cell fluorescence microscopy over time. In infected cultures, translocation was determined in N+ cells. Data from 1500 cells/condition are shown. (**d**) *IFNB* and *CXCL10* expression in cells from (a) over time, n=3. (**e-i**) Primary bronchial human airway epithelial cells (HAEs) were infected with 1500 E copies/cell of the indicated variants in the presence or absence of 5μM ruxolitinib, n=3. Viral replication was measured by (**e**) viral release into apical washes over time. Replication measured by apical release in HAEs infected with (**f**) Alpha WT or (**g**) ΔOrf6 in the presence or absence of 5μM ruxolitinib. (**h**) *IFNB* and *CXCL10* expression in cells from (e). (**i**) Replication of RG viruses BA.5 WT, ΔOrf6 and Orf6 D61L isolates in Calu-3 cells infected with 2000 E copies/cell over time. (**j**) Western blot of RG virus infections in Calu-3 cells at 24hpi for spike, N, Orf6 and β-Actin. (**k**) *IFNB* expression in cells from (i) over time. (**l**) Expression of *IFNB*, *CXCL10* and *IFIT1* in Calu-3 cells at 24hpi with 2000 E copies/cell of the indicated viruses. (**m**) Western blot of STAT1-pY701, STAT1-pS727, total STAT1, IRF3-pS396, total IRF3, and β-Actin at 24hpi. Quantification of four independent western blots showing (**n**) IRF3-pS396, (**o**) total IRF3, (**p**) STAT1-pS727, (**q**) STAT1-pY701 and (**r**) total STAT1 over β-Actin at 24hpi. Quantification of (**s**) IRF3 and (**t**) STAT1 translocation detected by single-cell fluorescence microscopy over time. In infected cultures, translocation was determined in N+ cells. Data from 1500 cells/condition are shown. (**u, v**) Replication of BA.5 WT and ΔOrf6 in HAEs infected with 1500 E copies/cell in the (**u**) absence or (**v**) presence of 5μM ruxolitinib. Quantification of (**w**) IRF3 and (**x**) STAT1 translocation detected by single-cell fluorescence microscopy over time in Calu-3 cells stimulated with poly(I:C) or vehicle Lipofectamine2000 (L2K). Data from 1500 cells/condition are shown. For indicated statistical comparisons at each time point in a, d-I, m, o-s, v and w one-way ANOVA with Dunnett’s post-test was used. In c, x and y, groups were compared by Kruskal-Wallis test at each time point. For j and l, RG mutants were compared to WT using a two-way ANOVA and Bonferroni post-test. Colors indicate comparator (BA.5 ΔOrf6, purple; BA.5 Orf6 D61L, light pink). Mean+/-SEM or individual datapoints are shown. hpi, hours post infection. *, p<0.05; **, p<0.01; ***, p<0.001; n.s., not significant.

Orf6 is a multifunctional viral accessory protein that modulates expression of host and viral proteins^33,34^. Orf6 selectively inhibits host nuclear transport to potently antagonize antiviral responses during infection. To probe Orf6 mechanisms, and its contribution to enhanced antagonism by the VOCs, we used reverse genetics to introduce two stop-codons into the Orf6 coding-sequence of both Alpha (Alpha ΔOrf6) and BA.5 (BA.5 ΔOrf6), which we confirmed abolished Orf6 expression during infection (Fig. 3b,j). While Alpha ΔOrf6 replicated similarly to parental wild type (WT) virus up to 24hpi (Fig. 3a,b), we observed enhanced *IFNB* and *CXCL10* expression (Fig. 3d) and protein secretion (Extended Data Fig. 5b) during Alpha ΔOrf6 infection of Calu-3 cells compared to WT virus. Moreover, increased IRF3 nuclear translocation was evident after Alpha ΔOrf6 infection at 24hpi using single cell quantitative immunofluorescence microscopy (Fig. 3c and Extended Data Fig. 5a). This suggests an important role for Orf6 in innate immune antagonism^16,34,35^ and is consistent with suppression of IRF3 nuclear transport in Orf6 overexpression studies^34–36^. The reduction in Alpha ΔOrf6 replication at 48hpi, and N and spike protein expression at 24hpi, that was rescued by ruxolitinib treatment, is also consistent with greater IFN mediated suppression of the Orf6 deletion mutant (Fig. 3b and Extended Data Fig. 5c).

Alpha ΔOrf6 also replicated less well than WT in HAE cells (Fig. 3e-g and Extended Data Fig. 5d). *IFNB* and *CXCL10* gene induction, normalized to *GAPDH*, were similar after Alpha ΔOrf6 and WT infection (Fig. 3h), despite lower E RNA levels for Alpha ΔOrf6, consistent with increased innate immune induction by the deletion virus. Importantly, Alpha ΔOrf6 was more sensitive to ruxolitinib treatment than WT, consistent with the notion that increased IFN induction caused reduced replication of Alpha ΔOrf6 (Fig. 3f,g). To address the role of Orf6 during BA.5 infection, we compared replication of a BA.5 ΔOrf6 mutant with parental BA.5 WT virus. We also generated the first example of a BA.5 mutant bearing the Orf6 D61L mutation found in BA.2 and BA.4, that has been proposed to reduce Orf6 function^25,31^ (Fig. 3i,j). Consistent with SARS-CoV-2 Alpha ΔOrf6 results, BA.5 ΔOrf6 showed a replication defect at 48hpi compared to BA.5 WT, and triggered significantly enhanced innate immune responses evidenced by enhanced *IFNB* and ISG induction (Fig. 3i-l). Deletion of Orf6 in BA.5 also increased the degree of infection-induced IRF3 and STAT1 phosphorylation (Fig. 3m-r) and nuclear translocation (Fig. 3s,t). This demonstrates that Orf6 loss enhances IRF3 and STAT1 activation despite similar levels of infection, confirming the important role of Orf6 in innate immune suppression and in distinguishing BA.5 from earlier Omicron subvariants. Infection of HAEs confirmed reduced viral replication of the BA.5 ΔOrf6 compared to WT BA.5, while viral release remained comparable (Fig 3u,v and Extended Data Fig. 5e). ISG expression in HAEs was similar between WT and mutant despite lower E RNA levels during BA.5 ΔOrf6 infection, suggesting greater induction of innate immunity in the absence of Orf6 in these cells (Extended Data Fig. 5f). Interestingly, introducing the C-terminal D61L mutation into BA.5 Orf6 resulted in an intermediate innate immune phenotype measured by increased induction of *IFNB*, *CXCL10* and *IFIT1* expression by the mutant virus (Fig. 3l). IRF3-phosphorylation and translocation were equivalent between BA.5 WT and Orf6 D61L (Fig 3n-s), whereas STAT1 translocation was not antagonized by Orf6 D61L (Fig. 3t), in line with reports of a partial loss of Orf6-function in the D61L mutation^25,31^. These data suggest complex adaptation of Orf6 manipulation of innate immunity during SARS-CoV-2 Omicron lineage adaptation.

During the course of this study, SARS-CoV-2 has continued to evolve and produce new Omicron subvariants (Fig. 4a and Extended Data Fig. 6a). Omicron subvariants BA.2.75, XBB.1, XBB.1.5 and BQ.1.1 have acquired increased ACE2 binding and enhanced adaptive immune evasion^37,38–40^. To test whether enhanced innate immune antagonism is consistently associated with globally successful subvariants, we compared BA.2.75, XBB.1, XBB.1.5 and BQ.1.1 isolates to BA.2 and BA.5 (Fig. 4). We equalized virus dose by Nsp12 RNA copies (RT-qPCR), a measurement of genomic RNA, rather than E RNA copies, due to accumulation of mutations in the E gene of later Omicron subvariants, including in the region detected by our RT-qPCR assay. We found that all Omicron subvariants retained an enhanced dependence on cathepsin, here measured in A549 cells expressing ACE2 and TMPRSS2 (Extended Data Fig. 6b). BA.2.75, XBB.1 (two independent isolates) and XBB.1.5, derived from the parental BA.2 lineage^38,40^, replicated comparably to earlier BA.2 and BA.5 in Calu-3 and HAEs (Fig. 4b-e and Extended Data Fig. 6c-i). BQ.1.1, which has arisen from BA.5^40^, displayed some reduction of replication in epithelial cells (Fig. 4d,e,h and Extended Data Fig. 6e,f,i). Similar to BA.5, we found that all subsequent Omicron subvariants tested triggered significantly less *IFNB* and *CXCL10* expression than BA.2 at 24hpi (Fig. 4f,g). All Omicron subvariants derived from BA.2 (BA.2.75, XBB.1 and XBB.1.5) showed reduced rescue by ruxolitinib treatment, as well as reduced induction of or sensitivity to IFN, similar to BA.5 (Fig. 4h and Extended Data Fig. 6f). Strikingly, like BA.5, enhanced innate immune evasion by these more recent subvariants was accompanied by increased Orf6 expression for the majority of isolates (Fig. 4i,j). Reduced BQ.1.1 replication in Calu-3 cells (Fig. 4d and Extended Data Fig. 6e) prevented Orf6 and N detection in the absence of ruxolitinib (Fig. 4i). Reduced innate activation by recent Omicron subvariants also correlated with reduced IRF3-phosphorylation compared to BA.2, and reduction of STAT1 serine-phosphorylation was principally observed for XBB.1 and XBB.1.5 variants (Fig. 4k-m and Extended Data Fig. 6j-l). Together these data are consistent with a trend for ongoing Omicron evolution enhancing Orf6 expression as it adapts to the human population leading to reduced innate immune responses, detectable at the level of IFN and ISG expression, and at the level of transcription factor phosphorylation and nuclear translocation. This study considering Omicron variants is very reminiscent of our previous observation of enhanced expression of key innate immune antagonists Orf6, N and Orf9b in VOCs Alpha to Delta suggesting a common evolutionary trajectory to combating human innate immunity and enhancing transmission^16,25^.

**Fig. 4.**
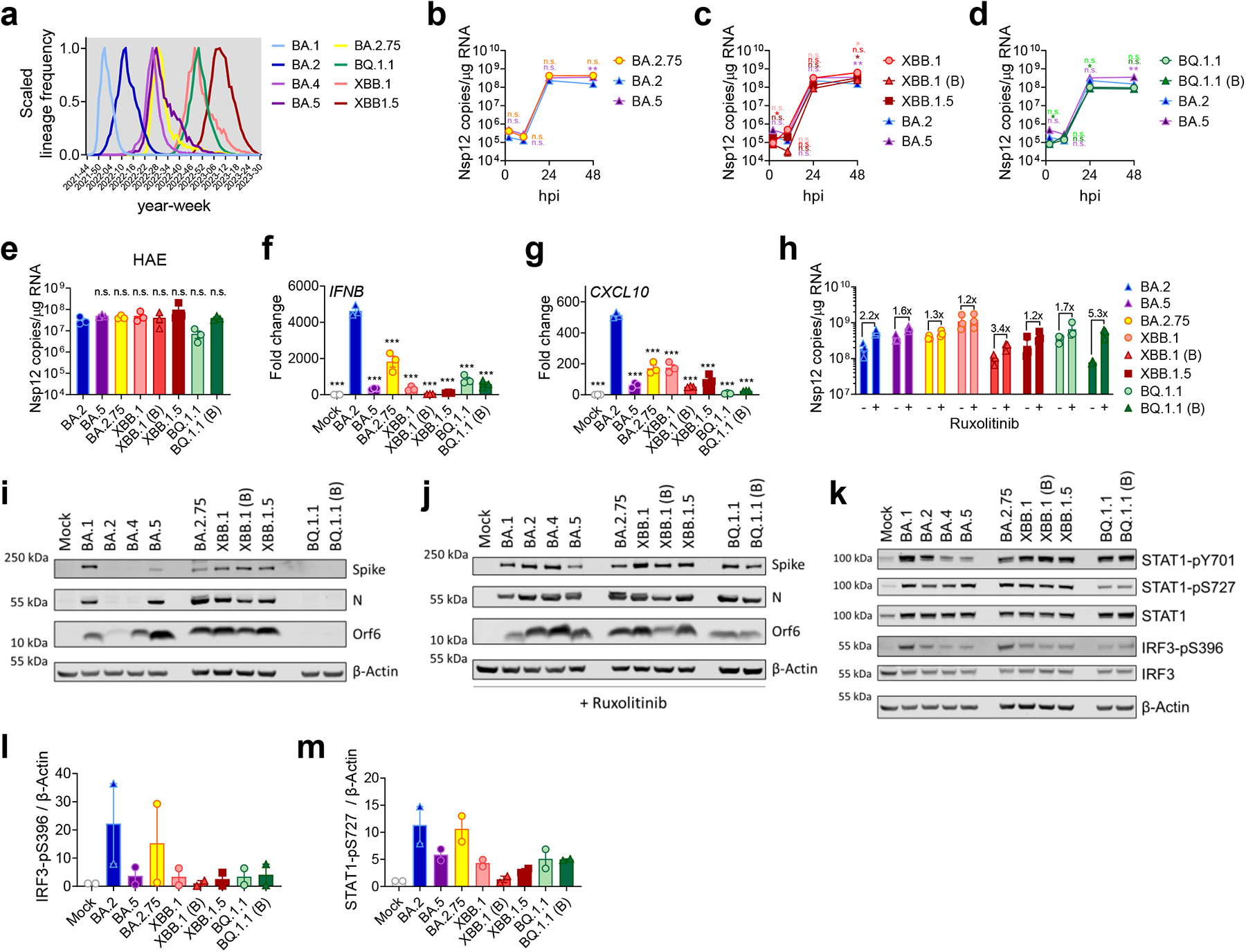
Innate immune phenotype of dominant Omicron subvariants. (**a**) Global SARS-CoV-2 variant sequence counts over time (scaled per variant), extracted from CoV-Spectrum using genomic data from GISAID. (**b-d**) Calu-3 cells were infected with 2000 Nsp12 copies/cell. Replication of Omicron subvariants compared to BA.2 (blue) and BA.5 (purple) measured by Nsp12 copies/μg RNA is shown for (**b**) BA.2.75 (yellow; Ο), (**c**) XBB-subvariants (XBB.1: light red, Ο; XBB.1 (B): red, Δ; XBB.1.5: dark red, □) and (**d**) BQ.1.1 (BQ.1.1: light green, Ο; BQ.1.1 (B): dark green, Δ) isolates. (**e**) HAEs were infected with 1500 Nsp12 copies/cell and intracellular Nsp12 copies measured at 72hpi. (**f**) *IFNB* and (**g**) *CXCL10* expression in Calu-3 cells infected with 2000 Nsp12 copies/cell of the indicated Omicron subvariants at 24hpi. (**h**) Viral replication of indicated variants in Calu-3 cells in the presence or absence of 5μM ruxolitinib at 48hpi. Numbers indicate fold change in replication in the presence of 5μM ruxolitinib. (**i,j**) Western blot of Orf6, N, spike and β-Actin at 48hpi in infected Calu-3 cells from (b-d) in the (**i**) absence or (**j**) presence of 5μM ruxolitinib. (**k**) Western blot of STAT1-pY701, STAT1-pS727, total STAT1, IRF3-pS396, total IRF3, and β-Actin in Calu-3 cells at 24hpi. Quantification of two independent western blots showing (**l**) IRF3-pS396 and (**m**) STAT1-pS727 over β-Actin at 24hpi. For b-d, variant replication was compared to BA.2 at each time point using a two-way ANOVA and Bonferroni post-test. Colors indicate comparator (BA.5, purple; BA.2.75, yellow; XBB.1, light red; XBB.1 (B), red; XBB.1.5, dark red; BQ.1.1, light green; BQ.1.1 (B), dark green). For e-g, one-way ANOVA with Dunnett’s post-test was used to compare all variants to BA.2. Mean+/-SEM or individual datapoints are shown. hpi, hours post infection. *, p<0.05; **, p<0.01; ***, p<0.001; n.s., not significant.

We propose a model in which the earliest host innate immune responses make an important contribution to SARS-CoV-2 transmission by influencing whether interactions with the first few cells in the airway establish a productive infection. In this model, viruses with enhanced ability to evade or antagonize innate immunity, for example through increased Orf6 and N expression, will transmit with greater frequency because they are better at avoiding inducing, or better at shutting down host responses that suppress this earliest replication.

How early viral manipulation of the host innate immune response influences disease is less clear. We hypothesize that once infection of the airway is irrevocably established, innate immune suppression that permits greater levels of viral replication may in turn lead to increased disease, simply due to greater viral burden and greater inflammatory responses. This model is supported by longitudinal nasal sampling of SARS-CoV-2 infected patients shortly after confirmation of infection, which revealed pronounced and early upregulation of an innate immune response in epithelial cells that rapidly declines after symptom onset ^41^. Concordantly, higher baseline antiviral gene expression and more potent innate induction in the nasal epithelium of children are associated with less severe infection outcomes compared to adults^42^. Like others, we assume this is explained by reduced viral loads reducing disease and early IFN protecting against transmission, with late IFN responses contributing to disease^43^. Similarly, inborn errors of innate antiviral mechanisms and IFN autoantibodies are associated with severe COVID19^44–48^, assumed to be explained by greater viral loads driving increased inflammatory disease. Furthermore, clinical trials of JAK/STAT inhibitors reduced COVID-19 mortality after hospitalization^49^. Considering an unrelated virus, simian immunodeficiency virus (SIV) in macaques, may be relevant. Here transmission efficiency and subsequent disease are also influenced by IFN at the site of infection^50^. In all these examples, early IFN is beneficial, reducing transmission, but late IFN is bad, increasing symptoms. Human SARS-CoV-2 challenge studies are expected to help us understand the effect of these dynamics and innate immune contributions to transmission and disease by permitting sampling before exposure and during the earliest time points post infection with careful assessment of disease in a highly controlled environment^51,52^.

We have focused on changes in expression of N and Orf6 but we expect that other viral genes contribute to evasion of innate immunity and adaptation to humans. In contrast to common cold coronaviruses, SARS-CoV-2 and its relatives encode a broad range of accessory genes^53,54^ which antagonize innate immunity and likely contribute to effective transmission between species. Our data suggest that upregulation of Orf6 expression is a central feature of SARS-CoV-2 adaptation to humans. Our observations using Orf6-deletion viruses confirm Orf6 to be a potent viral innate immune antagonist and are consistent with a model in which, like Alpha, Omicron subvariant enhancement of Orf6 expression contributes to the reduced innate immune response to infection compared earlier Omicron viruses. Orf6 upregulation by BA.5 may, in part, explain increased pathogenicity *in vivo*^1,2^. This notion is supported by ΔOrf6 SARS-CoV-2 infection of transgenic mice or hamsters, where the Orf6 mutant causes less severe disease and there is quicker recovery from infection, despite comparable viral loads in nose and lungs^31,55^. Expression of accessory and structural proteins as subgenomic RNAs during SARS-CoV-2 replication provides an elegant mechanism to selectively regulate their abundance during adaptation to host, as the level of each sgRNA and thus protein can be independently adjusted by mutation, as we found for VOCs Alpha to Delta^16,25^.

The earliest Omicron subvariants BA.1 and BA.2 outcompeted Delta despite not enhancing innate immune antagonism, explained by extensive antibody escape and improved spike function/stability^56,57^. This suggests adaptive immunity was the strongest selection force for Omicron emergence and global dominance. We hypothesize that the acquisition of enhanced innate immune suppression by Omicron lineage variants after their initial emergence required selection for improved transmission and dominance. Thus, innate immune escape may be the second dominant selective force the virus experiences after escape from neutralizing antibodies in a population with pre-existing immunity from prior infection and vaccination. We propose that evolving to better manage host innate immunity for improved transmission is a central feature of species-specific host-adaptation for all emerging viruses. Intriguingly, SARS-CoV-2 continues to jump species barriers and has been detected infecting 34 different animal species so far (https://vis.csh.ac.at/sars-ani/), illustrating its remarkable capacity to universally antagonize species specific innate immune responses. SARS-CoV-2 will be a fantastic model to further dissect species barriers to zoonotic spillovers and understand how viruses adapt to new species.

We propose that adaptation in spike and beyond also contributes to enhanced replication in human cells^16,25^. This may be important for outpacing early innate responses during transmission particularly in environments with a mix of permissive and non-permissive cells such as the upper human airways in which ACE2 is only expressed on ciliated cells^58^. Indeed, we have found that SARS-CoV-2 replicates more slowly in primary HAE cultures than in Calu-3 cells and that HAEs better recapitulate VOC replication advantages^16,25^. Primary HAEs complement more tractable monoculture models, such as Calu-3 that allow mechanistic studies. We propose that linking VOC genotype to phenotype in multiple models will be essential for effective prediction of novel variant behavior. Moreover, understanding how adaptive changes in spike leading to altered viral tropism influence innate immune responses also warrants further study.

We propose that this study adds to the body of evidence for innate immunity as a key barrier which must be overcome by all pandemic zoonotic viruses, particularly in the absence of immune memory in an exposure-naive species. This has also been elegantly demonstrated recently for influenza virus (IVA) where avian, but not human IVA, is efficiently restricted by human BTN3A3^59^, which like MX1^60^, can be overcome by adaptation to the human host. Innate immune evasion has also been linked to the single pandemic HIV-1 lineage^61^. Our findings herein have broad implications for understanding zoonotic pathogen emergence because they reveal molecular details of how SARS-CoV-2 Omicron subvariants have achieved dominance, unexpectedly by increasing specific protein expression rather than adapting by protein coding mutation. Crucially, they suggest that improvements in innate immune evasion can continue to enhance transmission, even after establishment in humans, illustrating an inevitable ongoing trajectory of adaptation towards escape from the innate immune mechanisms that are the gatekeepers of transmission success.

## Figure legends

**Extended Data Fig. 1.**
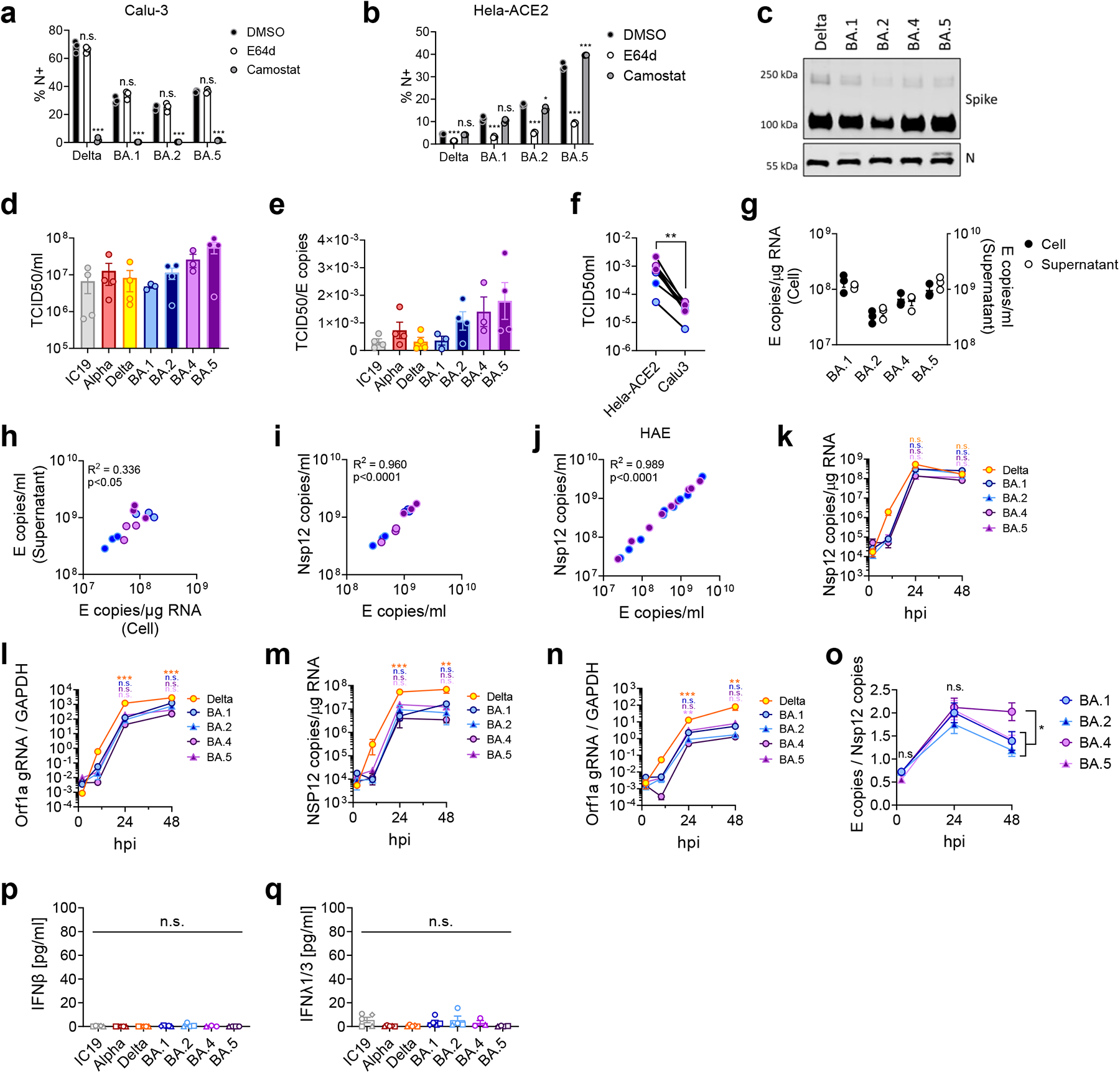
Replication measurements of SARS-CoV-2 variants. (**a**) Calu-3 and (**b**) Hela-ACE2 cells were infected with 1000 E copies/cell of the indicated variants in the presence of DMSO (-), 25μM E64d or 25μM camostat. Infection levels were measured at 24hpi by nucleocapsid expression (% N+ by flow cytometry), n=3. (**c**) Representative western blot of spike and N in purified SARS-CoV-2 virions, n=2. (**d**) Quantification of viral stock used in Fig. 1 and 2 by TCID50/ml on Hea-ACE2 cells. (**e**) Ratio of TCID50/ml over E copies/ml for virus stocks from (d). (**f**) TCID50/ml of indicated virus stocks measured on ACE2-Hela or Calu-3 cells. (**g-i**) Calu-3 cells were infected with 2000 E copies/cell of the indicated variants. At 24hpi, cells and culture supernatant were harvested to determine intracellular viral replication and virus release. (**g**) Intracellular replication (Cell) and viral release (Supernatant) was determined by quantification of E copies at 24hpi. (**h**) Correlation graph of intracellular E copies and virus released into supernatant at 24hpi. (**i**) Correlation of Nsp12 and E gene copies in supernatants from (g). (**j**) Correlation of Nsp12 and E copies in apical washes from HAEs infected with BA.2 (blue) or BA.5 (purple) (samples from Fig. 4). Calu-3 cells were infected with 2000 E copies/cell and viral replication measured by (**k**) Nsp12 copies/μg RNA or (**l**) Orf1a gRNA/*GAPDH* in cells from Fig. 1a. Viral replication measured by (**m**) Nsp12 copies/μg RNA or (**n**) Orf1a gRNA/*GAPDH* in cells infected with 200 E copies/ml from Extended Data Fig. 2h. (**o**) E copies/Nsp12 copies ratio in Calu-3 cells over time calculated from three independent experiments. (**p**) IFNβ and (**q**) IFNλ1/3 levels detected in SARS-CoV-2 variant inoculum prepared from virus stocks prepped in Caco-2 cells. For statistical comparison in a, b, k-q, one-way ANOVA with a Bonferroni post-test was used. For a, b, groups were compared to DMSO. For k-o, groups were compared to BA.2 and colors indicate comparator (Delta, yellow; BA.1, blue; BA.4, purple; BA.5, pink). For f, groups were compared by paired Student’s t-Test. R^2^ and p-values in h-j were calculated using simple linear regression. Mean+/-SEM or individual datapoints are shown. hpi, hours post infection. gRNA, genomic RNA. *, p<0.05; **, p<0.01; ***, p<0.001; n.s., not significant.

**Extended Data Fig. 2.**
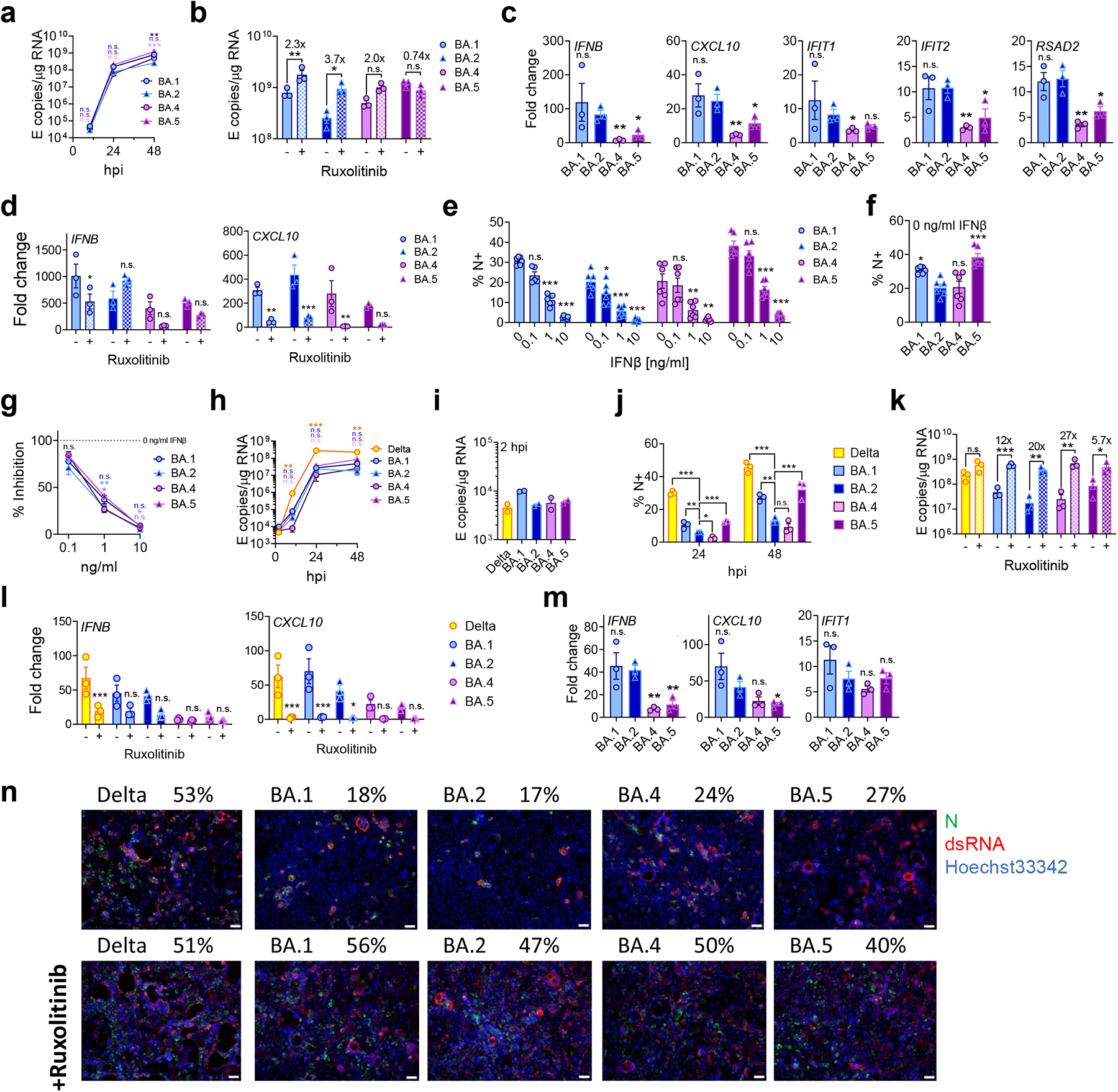
BA.5 displays enhanced innate immune antagonism during infection of airway epithelial cells. (**a-d**) Calu-3 infection with 2000 E copies/cell of BA.1 (blue; Ο), BA.2 (blue; Δ), BA.4 (purple; Ο) and BA.5 (purple; Δ), n=3. (**a**) Viral replication over time, n=3. (**b**) Viral replication of indicated variants in the presence or absence of 5μM ruxolitinib at 48hpi, n=3. (**c**) *IFNB*, *CXCL10*, *IFIT1, IFIT2* and *RSAD2* expression at 24hpi, n=3. (**d**) Expression of *IFNB* and *CXCL10* in the presence of ruxolitinib in cells from (c). (**e-g**) IFNβ-sensitivity of indicated variants during Calu-3 cell infection at 2000 E copies/cell. (**e**) Infection levels measured by % N+ at 24hpi at the indicated concentrations of IFNβ, n=6. (**f**) Infection levels in cells from (e) at 0ng/ml IFNβ, n=6. (**g**) Infection levels from (e) normalized to 0ng/ml IFNβ for each variant, n=6. (**h-m**) Calu-3 infection with 200 E copies/cell of Delta (yellow; Ο), BA.1 (blue; Ο), BA.2 (blue; Δ), BA.4 (purple; Ο) and BA.5 (purple; Δ), n=3. (**h**) Viral replication over time measured by RT-qPCR for intracellular E copies. (**i**) Viral E copies at 2hpi in cells from (h). (**j**) Infections levels measured by nucleocapsid expression (% N+ by flow cytometry). (**k**) Viral replication of indicated variants in Calu-3 cells from (a) in the presence or absence of 5μM ruxolitinib at 48hpi, n=3. (**l**) Expression of *IFNB* and *CXCL10* in the presence of ruxolitinib in cells from (h). (**m**) Expression of *IFNB, CXCL10* and *IFIT1* in infected cells at 24hpi in cells from (h). (**n**) Fluorescence microscopy of Calu-3 cells infected at 2000 E copies/cell at 48hpi in the presence or absence of 5μM ruxolitinib. Percentage infection quantified by dsRNA-positive cells is indicated per condition. Nucleocapsid, green; dsRNA, red; Hoechst33342, blue. Representative images shown. Scale bar, 50μm. For statistical comparisons, one-way ANOVA and Dunnett’s post-test were used. Groups were compared as indicated or with BA.2. For e, comparisons were made against 0ng/ml IFNβ for each variant. Colors in a, g and h indicate comparator (Delta, yellow; BA.1, blue; BA.4, purple; BA.5, pink). Mean+/-SEM or individual datapoints are shown. hpi, hours post infection. *, p<0.05; **, p<0.01; ***, p<0.001; n.s., not significant.

**Extended Data Fig. 3.**
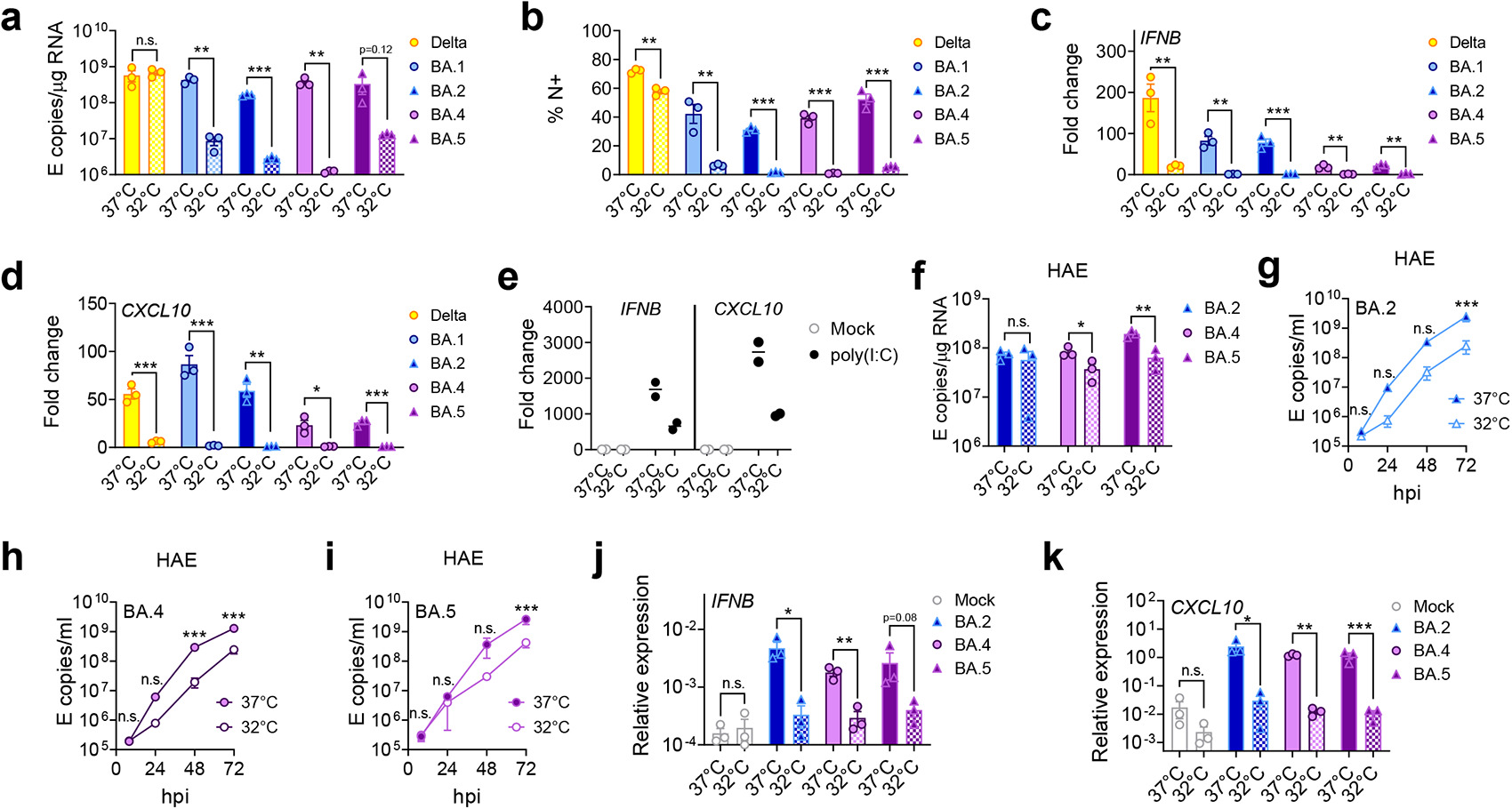
Entry and replication characteristics of Omicron subvariant BA.5. (**a-d**) Calu-3 cells were infected with 2000 E copies/cell at 37°C or 32°C, n=3. (**a**) Viral replication by RT-qPCR and (**b**) infection levels by flow cytometry at 24hpi. (**c**) *IFNB* and (**d**) *CXCL10* expression in cells from (a). (**e**) *IFNB* and *CXCL10* expression in response to poly(I:C) transfection in Calu-3 cells at 24h of stimulation, n=2. (**f-k**) Primary bronchial human airway epithelial cells (HAEs) were infected with 1500 E copies/cell of the indicated variants at 37°C or 32°C, n=3. Viral replication was measured by (**f**) intracellular E copies at 72hpi and viral release of (**g**) BA.2, (**h**) BA.4 and (**i**) BA.5 into apical washes over time. Relative expression of (**j**) *IFNB* and (**k**) *CXCL10* normalized to *GAPDH* in cells from (f). Fold changes are normalized to mock. Pairwise comparisons were performed using an unpaired two-tailed Student’s t-Test as indicated. For g-i, two-way ANOVA with a Bonferroni post test was used to compare temperatures at each time point. Mean+/-SEM or individual datapoints are shown. hpi, hours post infection. *, p<0.05; **, p<0.01; ***, p<0.001; n.s., not significant or exact p-value given (k).

**Extended Data Fig. 4.**
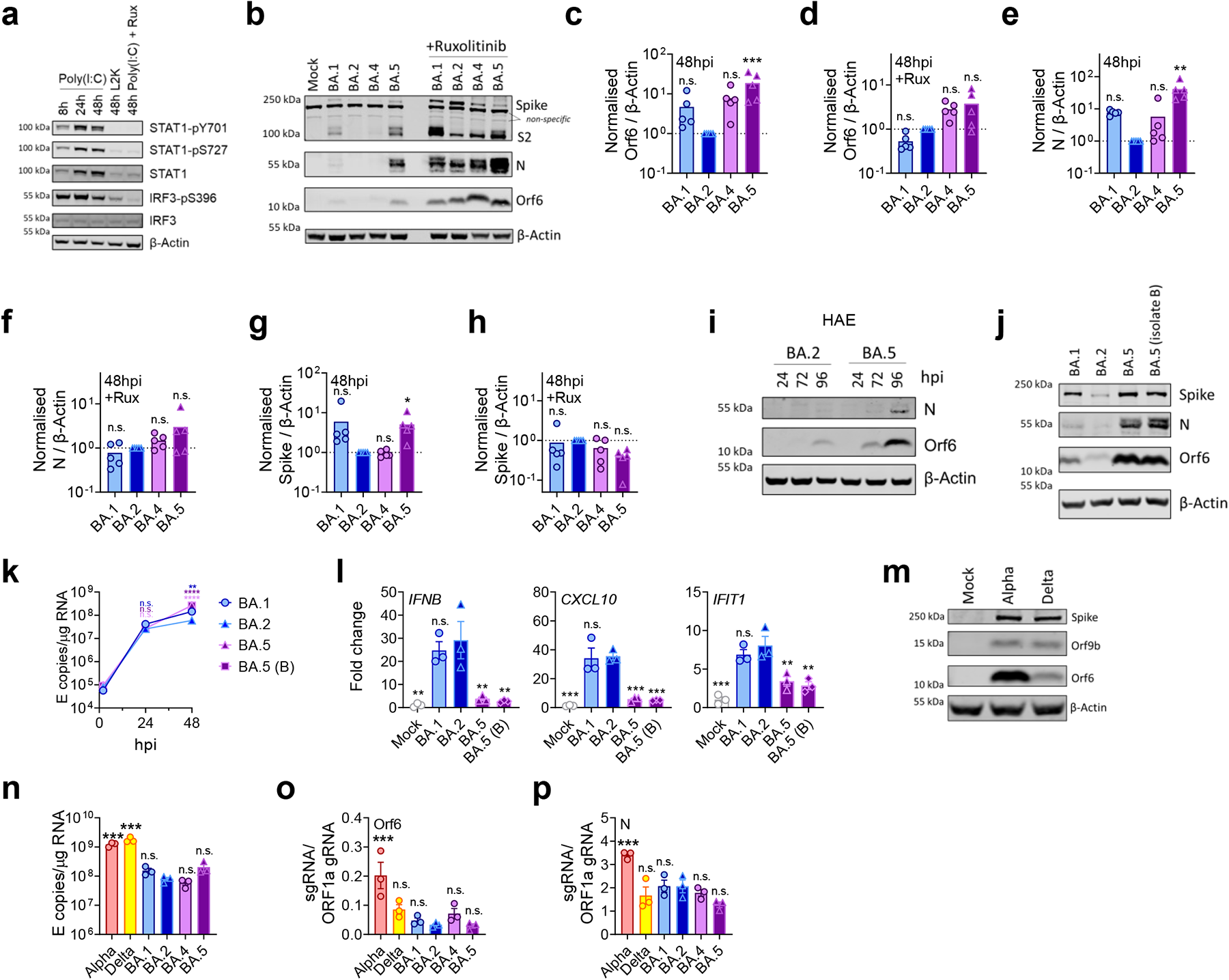
BA.5 efficiently expresses SARS-CoV-2 innate antagonists during airway epithelial cell infection. (**a**) Western blot of Calu-3 cells treated with poly(I:C), vehicle control lipofectamin2000 (L2K) or 5μM ruxolitinib where indicated. STAT1-pY701, STAT1-pS727, total STAT1, IRF3-pS396, total IRF3, and β-Actin are shown at indicated time points. (**b**) Western blot of Orf6, N, spike/S2 and β-Actin at 48hpi in infected Calu-3 cells +/-5μM ruxolitinib. Non-specific bands detected by polyclonal anti-spike primary antibody are indicated. (**c-h**) Quantification of viral protein expression from five independent western blots of infected Calu-3 cells at 48hpi +/-5μM ruxolitinib. (**c,d**) Orf6, (**e,f**) N and (**g**,**h**) spike were normalized to β-Actin over BA.2. (**i**) Western blot of Orf6 and N expression by HAEs infected with 1500 E copies/cell of BA.2 or BA.5 over time. (**j**) Western blot of Orf6, N, spike and β-Actin at 48hpi in Calu-3 cells infected with BA.1, BA.2 and two independent BA.5 isolates. (**k,l**) Calu-3 cells were infected with BA.1, BA.2 and two independent BA.5 isolates and (**k**) replication measured over time. (**l**) Expression of *IFNB*, *CXCL10* and *IFIT1* is shown at 24hpi in cells from (k). (**m**) Western blot of Orf9b, Orf6, spike and β-Actin at 24hpi in Calu-3 cells infected with the indicated variants at 2000 E copies/cell. (**n**) Viral replication in Calu-3 cells by RT-qPCR at 24hpi, n=3. (**o**) Orf6 and (**p**) N sgRNA expression in cells from (n), n=3. For c-h, l, n-o, one-way ANOVA with Dunnett’s post-test was used to compare BA.2 with other variants. For k, two-way ANOVA with a Bonferroni post-test was used to compare variants with BA.2 at each time point. Colors indicate comparator (BA.1, blue BA.5, purple; BA.5 (B), pink). Mean+/-SEM or individual datapoints are shown. hpi, hours post infection. sgRNA, subgenomic RNA. *, p<0.05; ***, p<0.001; n.s., not significant.

**Extended Data Fig. 5.**
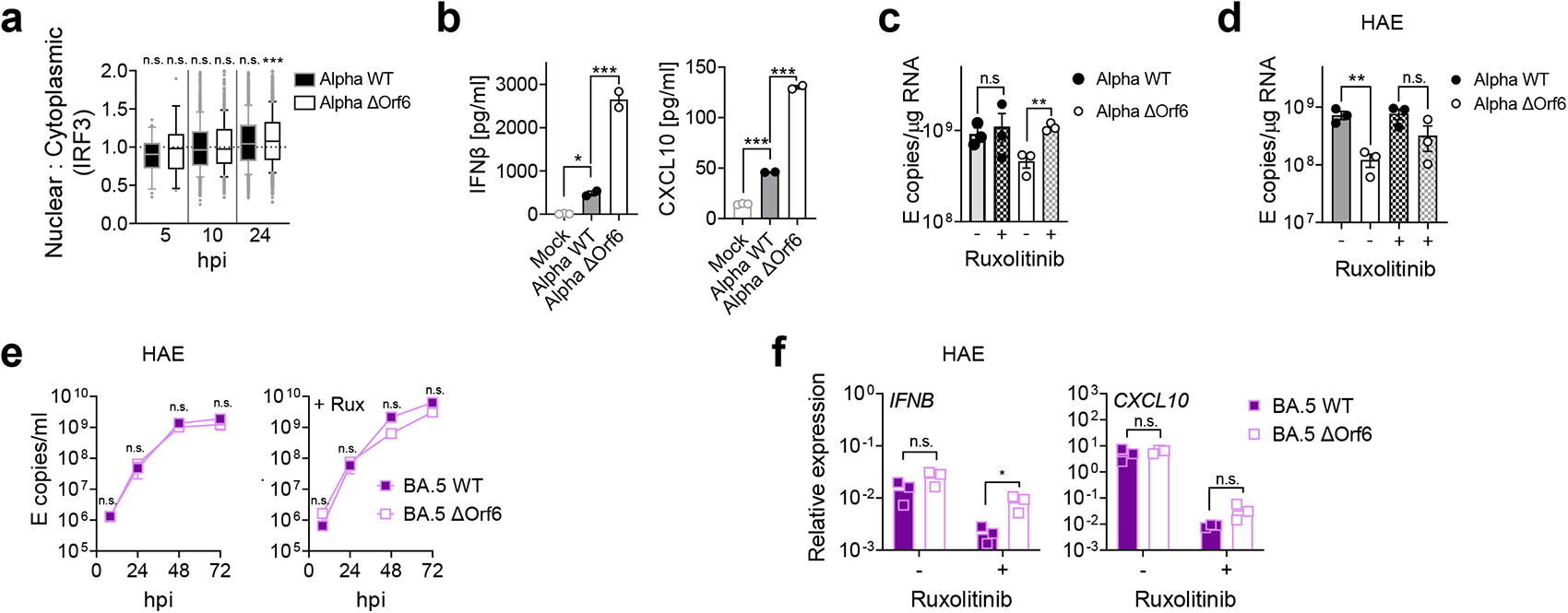
Orf6 expression is a major determinant of enhanced innate immune antagonism by emerging VOCs. (**a**) Quantification of IRF3 translocation in Calu-3 cells infected with Alpha WT and ΔOrf6 detected by single-cell fluorescence microscopy over time. Data from 1500 cells/condition are shown. (**b**) IFNβ and CXCL10 secretion from infected Calu-3 cells measured at 48hpi, n=2. (**c**) Viral replication in the presence or absence of 5μM ruxolitinib at 48hpi in cells from (Fig. 3a). (**d**) Primary bronchial human airway epithelial cells (HAEs) were infected with 1500 E copies/cell of the indicated variants in the presence or absence of 5μM ruxolitinib, Intracellular E copies are shown (n=3). Apical washes are shown in Fig. 3e-g. (**e, f**) Infection of HAEs with BA.5 WT or BA.5 ΔOrf6 with 1500 E copies/cell showing (**e**) viral release into apical washes over time or (**f**) *IFNB* and *CXCL10* expression at 72hpi. For a, Kruskal-Wallis test was used to compare groups to mock at each time point. In b-d, one-way ANOVA with Dunnett’s post-test was used to compare groups as indicated. For e, groups were compared at each time point using a two-way ANOVA with a Bonferroni post-test. Groups in f were compared by paired Student’s t-Test. *, p<0.05; **, p<0.01; ***, p<0.001; n.s., not significant.

**Extended Data Fig. 6.**
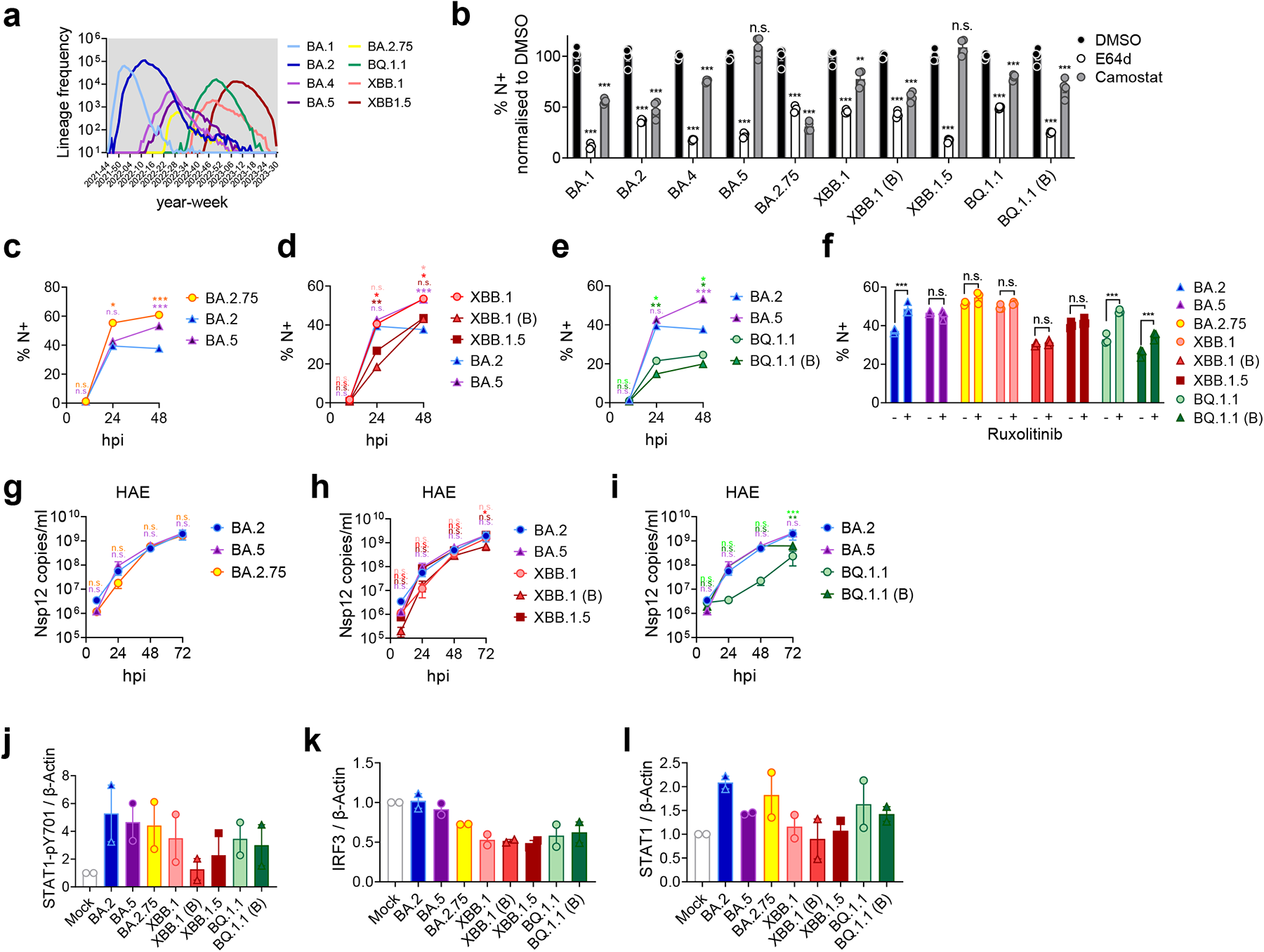
Innate immune phenotype of dominant Omicron subvariants. (**a**) Absolute global SARS-CoV-2 variant sequence counts over time, extracted from CoV-Spectrum using genomic data from GISAID. (**b**) ACE2/TMPRSS2-A549 cells were infected with 2000 Nsp12 copies/cell of the indicated SARS-CoV-2 variants in the presence of DMSO, 25μM E45d or 25μM camostat. Infection levels were determined by N-positivity at 24hpi. (**c-e**) SARS-CoV-2 Omicron subvariants infection of Calu-3 cells determined by N-positivity over time for the indicated subvariants in cells from Fig. 4b-d with (**c**) BA.2.75 (yellow; Ο), (**d**) XBB-subvariants (XBB.1: light red, Ο; XBB.1 (B): red, Δ; XBB.1.5: dark red, □) and (**e**) BQ.1.1 (BQ.1.1: light green, Ο; BQ.1.1 (B): dark green, Δ) isolates shown. (**f**) Infection levels of indicated variants in Calu-3 cells in the presence or absence of 5μM ruxolitinib at 48hpi in cells from Fig. 4h. (**g-i**) Primary bronchial human airway epithelial cells (HAEs) were infected with 1500 E copies/cell of the indicated variants. Viral replication was measured by viral release into apical washes over time in cells from Fig. 4e. (**g**) BA.2.75, (**h**) XBB-subvariants and (**i**) BQ.1.1 isolates are shown compared to BA.2 (blue) and BA.5 (purple). Quantification of two independent western blots showing (**j**) STAT1-pY701, (**k**) total IRF3 and (**l**) total STAT1 over β-Actin at 24hpi. For b, treatments were compared to DMSO for each variant using one-way ANOVA and Dunnett’s post-test. For c-i, variant infection levels were compared to BA.2 at each time point by two-way ANOVA and Bonferroni post-test. Colors indicate comparator (BA.5, purple; BA.2.75, yellow; XBB.1, light red; XBB.1 (B), red; XBB.1.5, dark red; BQ.1.1, light green; BQ.1.1 (B), dark green). Mean+/-SEM or individual datapoints are shown. hpi, hours post infection. *, p<0.05; **, p<0.01; ***, p<0.001; n.s., not significant.

**Extended Data Table 1.**
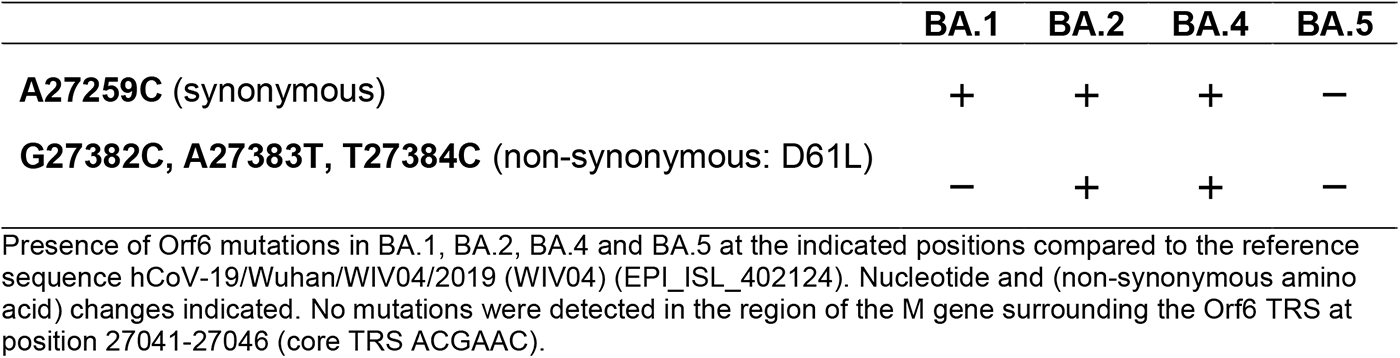
Orf6 mutations detected in the Omicron subvariants.

**Extended Data Table 2.** Nucleocapsid (N) mutations detected in the Omicron subvariants.

|  | BA.1 | BA.2 | BA.4 | BA.5 |
| --- | --- | --- | --- | --- |
| **C28311T** (non-synonymous: P13L) | + | + | + | + |
| **A28330G** (synonymous) | – | – | – | + |
| **A28363T** (synonymous) | + | + | + | + |
| **Deletion 28364-28372** (31-33del) | + | + | + | + |
| **C28724T** (non-synonymous: P151S) | – | – | + | – |
| **G28881A, G28882A** (non-synonymous R203K<sup>†</sup>) | + | + | + | + |
| **G28883C** (non-synonymous G204R<sup>†</sup>) | + | + | + | + |
| **A29510C** (non-synonymous S413R) | – | + | + | + |

Presence of N mutations in BA.1, BA.2, BA.4 and BA.5 at the indicated positions compared to the reference sequence hCoV-19/Wuhan/WIV04/2019 (WIV04) (EPI\_ISL\_402124). Nucleotide and (non-synonymous amino acid) changes indicated. BA. 1, BA.2, BA.4 and BA.5 carry nucleotide substitution A28271T, changing their Kozak initiation context from adequate (A in –3, T in +4) to the weak (T in –3, T in +4) as previously described<sup>16</sup>.
<sup>†</sup> The non-synonymous mutations R203K-G204R confer a partial TRS for N\* sgRNA<sup>16</sup>.

## Methods

### Cell culture

Calu-3 cells were purchased from AddexBio (C0016001), Caco-2 cells were a kind gift from Dalan Bailey (Pirbright Institute), Hela-ACE2 cells were a gift from James E Voss^62^ and A459 cells expressing ACE2 and TMPRSS2 were a kind gift from Massimo Palmarini ^19^.Cell lines were cultured in Dulbecco’s modified Eagle Medium (DMEM) supplemented with 10% heat-inactivated FBS (Labtech) and 100U/ml penicillin/streptomycin. Cells were passaged at 80-90% confluence. For infections, Calu-3 and Caco-2 cells were seeded at 2x10^5^ cells/ml and Hela-ACE2 cells at 1x10^5^ cells/ml and grown to 60-80% confluence for experiments^15,16^. Primary normal (healthy) bronchial epithelial (NHBE-A) cells from two independent donors were cultured for five to seven passages and differentiated at an air-liquid interface as previously described^15^. After 21-24 days of differentiation, cells were used in infection experiments.

### Viruses

SARS-CoV-2 lineages Alpha (B.1.1.7), Delta (B.1.617.2)^18^ and Omicron (lineage B.1.1.529.1/BA.1, lineage B.1.1.529.2/BA.2, lineage BA.2.75 (BA.2.75.3) lineage BQ.1.1 (BQ.1.1.1), lineage XBB.1) isolates were a gift from Wendy Barclay (Imperial College London, UK). Omicron BA.4 (lineage B.1.1.529.4), BA.5 (lineage B.1.1.529.5), BQ.1.1 (B) (BQ.1.1.15) and lineage XBB.1.5 (XBB.1.5.13) were a gift from Alex Sigal and Khadija Khan (Africa Health Research Institute, Durban, South Africa)^5,13^. SARS-CoV-2 BA.5 (B) (SARS-CoV-2/Norway/20365/2022) was obtained from the Norwegian Institute of Public Health, Oslo, Norway. Omicron isolate identity was confirmed by full genome sequencing and assigned by Nextclade v.2.14.1 (https://clades.nextstrain.org)^63,64^ . Alpha Orf6 deletion virus (Alpha ΔOrf6) was achieved by mutation of the first two methionines: M1L (A27216T) and M19L (A27200T). Reverse genetics derived viruses were generated as previously described^65,66^. In brief, to generate the WT SARS-CoV-2 Alpha variant, a set of overlapping viral genomic cDNA fragments were chemically synthesized (GENEWIZ, Germany). The cDNA fragment representing the 5’ terminus of the viral genome contained the bacteriophage T7 RNA polymerase promoter and the fragment representing the 3’ terminus contained the T7 RNA polymerase termination sequences. These fragments were then assembled into a full-length Alpha cDNA genome using the Transformation-Associated Recombination (TAR) in yeast method^65^ . To generate the Alpha virus carrying the ATG codon changes (M1L and M19L) in its Orf6 gene (to generate Alpha ΔOrf6), the relevant cDNA fragments were chemically synthesized (Thermofisher, UK) and the mutant viral genome assembled using TAR in yeast as described above. We similarly generated WT BA.5, BA.5 ΔOrf6 (carrying M1L and M19L changes), and BA.5 Orf6 D61L (generated by introducing the GAT→CTC nucleotide change found in BA.2) using TAR in yeast except that the assembled cDNA genomes were placed under the control of the human cytomegalovirus promoter and the relevant termination sequences. The assembled WT and Orf6 null mutant genomes were transfected into BHK-hACE2-N cells stably expressing the SARS-CoV-2 N and the human ACE2 gene for virus rescue^67^. The rescued viruses were passaged once (P1 stock) in Vero.E6 cells and their full genomes sequenced using Oxford Nanopore as previously described ^68^. For Alpha and BA.5 the RG-derived viruses are referred to as WT, ΔOrf6 or Orf6 D61L to differentiate them from the clinically isolated viruses used in all other experiments. All viruses were propagated by infecting Caco-2 cells in DMEM culture medium supplemented with 1% FBS and 100U/ml penicillin/streptomycin at 37 °C as previously described^15,16^. Virus was collected at 72 hpi and clarified by centrifugation at 2,100xg for 15 min at 4°C to remove any cellular debris. Virus stocks were aliquoted and stored at −80 °C. Virus stocks were quantified by extracting RNA from 100 µl of supernatant with 1 µg/ml carrier RNA using Qiagen RNeasy clean-up RNA protocol, before measuring viral E RNA copies per ml by RT-qPCR^15,16^. For experiments including Omicron subvariants XBB.1 and BQ.1.1, stocks and viral replication were quantified using Nsp12 RNA copies due to accumulation of mutations in the E gene of these variants, including in the region detected by our RT-qPCR assay. Virus titres were determined by TCID50 in Hela-ACE2 cells. 10^4^ cells were seeded in 96-well plates in 100μl. The next day, seven 10-fold serial dilutions of each virus stock or supernatant were prepared and 50 µl added to the cells in quadruplicate. Cytopathic effect (CPE) was scored at 48-72hpi. TCID50 per ml was calculated using the Reed & Muench method, and an Excel spreadsheet created by B. D. Lindenbach was used for calculating TCID50 per ml values^69^.

To generate SARS-CoV-2 lineage frequency plots for BA.1, BA.2, BA.4, BA.5, BA.2.75, BQ.1.1, XBB.1 and XBB.1.5 (Fig. 4a and Extended Data Fig. 6a), respective sequence counts were extracted from CoV-Spectrum (cov-spectrum.org)^70^ using genomic data from GSAID^71^.

### Virus culture and infection

For infections, inoculum was calculated using E copies per cell quantified by RT-qPCR. Cells were inoculated with indicated variants for 2h at 37°C, subsequently washed with PBS and fresh DMEM culture medium supplemented with 1% FBS and 100U/ml penicillin/streptomycin was added. At the indicated time points, cells were collected for analysis. For primary HAE infections, virus was added to the apical side for 2-3 h at 37°C. Supernatant was then removed and cells were washed twice with PBS. All liquid was removed from the apical side and basal medium was replaced with fresh Pneumacult ALI medium for the duration of the experiment. Virus release was measured at the indicated time points by extracting viral RNA from apical PBS washes. For poly(I:C) (Peprotech) stimulations, cells were transfected with poly(I:C) using lipofectamine2000 (InvitroGen) in Opti-Mem (Thermo) for the indicated times. For IFN-sensitivity assays, cells were pre-treated with indicated concentrations or recombinant human IFNβ (Peprotech) for 18h before infection. Cytokines were maintained throughout the experiment. For inhibition assays, cells were pre-treated with 5 μM Ruxolitinib (Cambridge Bioscience), 25μM camostat (Apexbio), 25μM E64d (Focus Biomolecules) or DMSO control for 2-3h before SARS-CoV-2 infection. Inhibitors were maintained throughout the infection.

### RT-qPCR of host and viral gene expression in infected cells

Infected cells were lysed in RLT (Qiagen) supplemented with 0.1% beta-mercaptoethanol (Sigma). RNA extractions were performed according to the manufacturer’s instructions using RNeasy Micro Kits (Qiagen) including on-column DNAse I treatment (Qiagen). cDNA was synthesized using SuperScript IV (Thermo) with random hexamer primers (Thermo). RT-qPCR was performed using Fast SYBR Green Master Mix (Thermo) for host gene expression and subgenomic RNA expression or TaqMan Master mix (Thermo Fisher Scientific) for viral RNA quantification, and reactions were performed on the QuantStudio 5 Real-Time PCR systems (Thermo Fisher Scientific). Viral E RNA copies were determined as described previously^15,16^. Viral subgenomic RNAs were detected using the same forward primer against the leader sequence paired with a sgRNA specific reverse primer^16,72,73^. Using the 2−ΔΔCt method, sgRNA levels were normalized to GAPDH to account for differences in RNA loading and then normalized to the level of Orf1a gRNA quantified in the same way for each variant to account for differences in the level of infection. Host gene expression was determined using the 2−ΔΔCt method and normalized to GAPDH expression. The following probes and primers were used:

*GAPDH* fw: 5’-ACATCGCTCAGACACCATG-3’, rv: 5’-TGTAGTTGAGGTCAATGAAGGG-3’; *IFNB* fw: 5’-GCTTGGATTCCTACAAAGAAGCA-3’, rv: 5’-ATAGATGGTCAATGCGGCGTC-3’; *CXCL10* fw: 5’-TGGCATTCAAGGAGTACCTC-3’, rv: 5’-TTGTAGCAATGATCTCAACACG-3’; *IFIT1* fw: 5’-CCTCCTTGGGTTCGTCTACA-3’, rv: 5’-GGCTGATATCTGGGTGCCTA-3’; *IFIT2* fw: 5′-CAGCTGAGAATTGCACTGCAA-3′, rv: 5′-CGTAGGCTGCTCTCCAAGGA-3′; *MX1* fw: 5′-ATCCTGGGATTTTGGGGCTT-3′, rv: 5′-CCGCTTGTCGCTGGTGTCG-3′; *MX2* fw: 5’-CAGCCACCACCAGGAAAC-3’, rv 5’-TTCTGCTCGTACTGGCTGTACAG-3’, *RSAD2* fw: 5′-CTGTCCGCTGGAAAGTG-3′, rv: 5′-GCTTCTTCTACACCAACATCC-3′; *DDX58* fw: 5’-CTGGACCCTACCTACATCCTG-3’, rv: 5’-GGCATCCAAAAAGCCACGG-3’. SARS-CoV-2 E Sarbeco fw: 5′-CGTTAATAGTTAATAGCGTACTTCTTTTTC-3′; SARS-CoV-2 E Sarbeco Probe1: 5′-FAM-ACACTAGCCATCCTTACTGCGCTTCG-TAMRA-3′; SARS-CoV-2 E Sarbeco rv 5′-ATATTGCAGCAGTACGCACACA-3′; SARS-CoV-2 Nsp12 fw: 5’-GAGTGAAATGGTCATGTGTGG-3’; SARS-CoV-2 Nsp12 rv: 5’-CATTGGCCGTGACAGCTTGAC-3’; SARS-CoV-2 Nsp12 Probe: 5’-CTCATCAGGAGATGCCACAACTGCTTATGCTAATAG-3’;5′ Leader fw: 5’-ACCAACCAACTTTCGATCTCTTGT-3’; Orf1a rv: 5’-CCTCCACGGAGTCTCCAAAG-3’; Orf6 rv:GAGGTTTATGATGTAATCAAGATTC; N rv: 5’-CCAGTTGAATCTGAGGGTCCAC-3’; Orf3a rv: 5’-GCAGTAGCGCGAACAAAAT-3’. S rv: 5’-GTCAGGGTAATAAACACCACGTG-3’.

### Flow cytometry

Adherent cells were trypsinized and fixed in 4% formaldehyde prior to intracellular staining for SARS-CoV-2 nucleocapsid (N) protein. For N detection, cells were permeabilized for 15 min with Intracellular Staining Perm Wash Buffer (BioLegend) and subsequently incubated with 1μg/ml CR3009 SARS-CoV-2 cross-reactive antibody (a gift from Laura McCoy) for 30 min at room temperature. Primary antibodies were detected by incubation with secondary Alexa Fluor 488-Donkey-anti-Human IgG (Jackson Labs). All samples were acquired on a BD Fortessa X20 or LSR II using BD FACSDiva software. Data was analyzed using FlowJo v10.6.2 (Tree Star).

### Cytokine secretion

Secreted mediators were detected in cell culture supernatants by ELISA. IFNβ, IFNλ1/3 and CXCL10 were measured using Human IFN-beta Quantikine ELISA Kit, Human IL-29/IL-28B (IFN-lambda 1/3) DuoSet ELISA or Human CXCL10/IP-10 DuoSet ELISA reagents (biotechne R&D systems) according to the manufacturer’s instructions.

### Western blotting

For detection of N, Orf6, Orf9b, spike and β-actin expression, whole-cell protein lysates were extracted with RIPA buffer, and then separated by SDS–PAGE, transferred onto nitrocellulose and blocked in PBS with 0.05% Tween 20 and 5% skimmed milk. Membranes were probed with rabbit-anti-SARS spike (Invitrogen, PA1-411-1165), mouse-anti-SARS-CoV-2 spike (GeneTex 1A9), rabbit-anti-Orf6 (Abnova, PAB31757), rabbit-anti-Orf9b (ProSci, 9191), Cr3009 SARS-CoV cross-reactive human-anti-N antibody (a gift from Laura McCoy, UCL), rabbit-anti-phospho-STAT1 (Ser727) (CellSignaling, Cat # 9177), rabbit-anti-phospho STAT1 (Tyr 701) (CellSignaling, Cat# 9167, clone 58D6), rabbit-anti-STAT1 (CellSignaling, Cat# 9172), anit-rabbit-IRF3 (CellSignaling, Cat# 4302), rabbit-anti-phospho IRF3 (CellSignaling, Cat# 29047, clone D6O1M) and rabbit-anti-beta-actin (SIGMA), followed by IRDye 800CW or 680RD secondary antibodies (Abcam, goat anti-rabbit, goat anti-mouse or goat anti-human). Blots were imaged using an Odyssey Infrared Imager (LI-COR Biosciences) and analyzed with Image Studio Lite software. For virion blots, live virus normalized by equal total E copies was purified across a 25% sucrose cushion and concentrated by centrifugation (2h 16500xg, 4^0^C).

### Immunofluorescence staining and image analysis

Infected cells were fixed using 4% PFA/formaldehyde for 1h at room temperature and subsequently washed with PBS. A blocking step was carried out for 35h at room temperature with 10% goat serum/1%BSA/0.001 Triton-TX100 in PBS. dsRNA and nucleocapsid detection were performed by primary incubation with rabbit-anti-IRF3 antibody (sc-33641, Santa Cruz), rabbit-anti-STAT-1 (D1K9Y, Abcam), mouse-anti-dsRNA (MABE1134, Millipore) and Cr3009 SARS-CoV cross-reactive human-anti-N antibodies for 18h and washed thoroughly in PBS. Primary antibodies detection occurred using secondary anti-rabbit-AlexaFluor-488. anti-mouse-AlexaFluor-568 and anti-human-Alexa647 conjugates (Jackson ImmunoResearch) for 1h. All cells were labeled with Hoechst33342 (H3570, Thermo Fisher). Images were acquired using the WiScan® Hermes 7-Colour High-Content Imaging System (IDEA Bio-Medical, Rehovot, Israel) at magnification 10X/0.4NA. Four channel automated acquisition was carried out sequentially. Images were acquired across a well area density resulting in 31 FOV/well and ∼20,000 cells.

Images were pre-processed by applying a batch rolling ball background correction in FIJI ImageJ software package^74^ prior to quantification. IRF3 and STAT1 translocation analysis was carried out using the Athena Image analysis software (IDEA Bio-Medical, Rehovot, Israel) and data post-processed in Python. For dsRNA, Infected cell populations were determined by thresholding of populations with greater than 2 segmented dsRNA punctae. For transcription factor translocation analysis, infected populations were determined by presence of segmented nucleocapsid signal within the cell.

### Statistical analysis

Statistical analysis was performed using GraphPad Prism9 and details of statistical test used are indicated. Data shows mean +/-SEM with significant differences or exact p-values indicated in the figures. Significance levels were defined as follows: *, p < 0.05; **, p < 0.01 and ***, p < 0.001.

## Data availability

All data generated or analyzed during this study are included in this manuscript (and its supplementary information files). No new algorithms were developed for this project.

## Acknowledgements

This work was funded by Wellcome Investigator Awards 223065 to C.J. and 220863 to G.J.T. G.J.T. and C.J. were also funded by MRC/UKRI G2P-UK National Virology consortium (MR/W005611/1) and the UCL COVID-19 fund. R.R is supported by a Marie Skłodowska-Curie Individual Fellowships no. 896014. P.B received funding from the European Research Council (ERC-Stg no. 639429), the Rosetrees Trust (M362-F1; M553), the NIHR GOSH BRC and the CF Trust (SRC006; SRC020). M.V.X.W. is supported by the NIHR Biomedical Research Centre at UCLH and IDEA Bio-Medical. The work at the CVR was funded by G2P-UK National Virology Consortium funded by the MRC (MR/W005611/1) and MRC grants (MC_UU12014/2 and MC_UU_00034/9) and the Wellcome Trust (206369/Z/17/Z). We are grateful to Wendy Barclay (Imperial College London, UK), Alex Sigal and Khadija Khan (AHRI, South Africa) and the Norwegian Institute of Public Health for provision of virus isolates and Dalan Bailey, Laura McCoy and James Voss for reagents. For the purpose of Open Access, the authors have applied a CC BY public copyright license to any Author Accepted Manuscript version arising from this submission.

## Author contributions

Conceptualisation: A-K.R., L.G.T., G.J.T. and C.J.

Designed experiments: A-K.R, L.G.T. V.C., W.F., G.d.L., G.J.T. and C.J.

Performed experiments: A-K.R, L.G.T., M.X.V.W., D.M., G.D., N.B., V.C., W.F., and G.d.L

Analyzed data: A-K.R., L.G.T., M.X.V.W., G.J.T. and C.J.

Generated and provided reagents: R.R. W.F., G.de L., V.M.C., J.T., P.B., M.P. and A.H.P.

Manuscript preparation and editing: A-K.R, L.G.T., D.M., A.H.P., M.P., G.J.T. and C.J.

Coordination and supervision: P.B., M.P., A.H.P., G.J.T. and C.J.

## Competing interests

The authors declare that they have no competing interests.

Correspondence and request for materials should be addressed to Ann-Kathrin Reuschl, Clare Jolly or Greg J. Towers.

